# Acoustic Masking of Electric Stimulation at Basal and Extra-Cochlear Sites

**DOI:** 10.64898/2026.09.16.752065

**Authors:** Patrick Hinz, Waldo Nogueira

## Abstract

Cochlear implant (CI) candidates with residual low-frequency hearing increasingly receive hearing-preserving electrode arrays to enable electric–acoustic stimulation (EAS). However, assessing very low-frequency hearing in newborns and young children remains difficult. Previous work has shown that acoustic stimulation can mask electrically evoked auditory percepts when stimulation sites are spatially separated. Whether such masking also occurs when electric stimulation is delivered near the round window (RW), prior to cochlear insertion, remains unclear.

This study investigated ipsilateral acoustic masking of electric stimulation delivered at apical, basal, and near-RW sites in 12 EAS users with residual low-frequency hearing implanted with short or partially inserted electrode arrays. Using a psychophysical paradigm, threshold elevations of electric pulse trains were measured during simultaneous acoustic stimulation. Electric stimulation was applied at extra-cochlear RW locations and intra-cochlear basal and apical electrodes, while stimulation parameters were systematically varied.

Electric stimulation near the RW reliably evoked auditory percepts without side effects, though optimal stimulation parameters and masking strength varied across participants. Strong acoustic masking was consistently observed for apical stimulation, whereas weaker but measurable masking occurred for basal and RW stimulation. A cochlear lumped-parameter model indicated that approximately 5% of current delivered at the RW can reach apical regions, explaining the observed masking by low-frequency acoustic tones. Masking strength correlated with residual hearing for apical stimulation only.

These findings indicate that acoustic masking of apical electric stimulation, which correlated with residual hearing, holds potential as a diagnostic tool for assessing low-frequency hearing post-operatively. The detection of masking during both RW and basal stimulation represents a promising result, indicating that these stimulation paradigms can produce quantifiable effects. Given the considerable variability observed among participants, further investigation is required to establish the extent to which masking measures reflect residual hearing or hearing loss. Nonetheless, these results provide novel insight into auditory perception and side effects associated with basal and extra-cochlear electric stimulation, informing future efforts to optimize this approach for low-frequency hearing assessment.

## 1. Introduction

Cochlear implants (CIs) have proven highly effective in restoring access to sound, particularly for speech perception in quiet environments, for individuals with profound sensorineural hearing loss. By bypassing damaged hair cells, CIs deliver electrical pulses via electrodes implanted in the cochlea, directly stimulating the auditory nerve. Recent advancements in surgical techniques and electrode array designs have enabled the preservation of residual low-frequency acoustic hearing during and after CI insertion in the same ear (Gantz and Turner, 2003; Gstoettner et al., 2006; von Ilberg et al., 1999). As a result, CI candidacy has expanded to include individuals with partial high-frequency hearing loss while maintaining low-frequency acoustic hearing in the apical region of the cochlea (von Ilberg et al., 2011). These patients benefit from electric-acoustic stimulation (EAS), where electrical stimulation of high-frequency sounds is delivered via the CI, while low-frequency sounds are acoustically amplified using a hearing aid or acoustic component. Initially, this was achieved using specially designed short electrode arrays that were fully inserted into the cochlea, covering only its basal portion in order to preserve residual low-frequency hearing at the apex (Lenarz et al., 2009, 2013). While such short arrays are associated with high hearing preservation rates, they provide only a limited cochlear coverage. Consequently, if residual hearing is subsequently lost, patients who must rely exclusively on electrical stimulation (ES-only) generally achieve significantly poorer speech understanding than recipients implanted with longer electrode arrays (Büchner et al., 2017; Lenarz et al., 2019). This trade-off has influenced contemporary surgical strategies. Rather than routinely fully inserting long electrode arrays, current practice aims to match electrode length, and consequently insertion depth, to the amount of residual hearing present in each candidate, since deeper insertion increases the risk of trauma to structures supporting residual hearing (Lenarz et al., 2019). Electrode arrays of varying lengths (e.g., 20, 24, and 28 mm) allow for this individualized selection. Where no sufficiently short array is available, or where the required coverage cannot otherwise be achieved, full-length arrays can instead be partially inserted, such that the most basal contacts remain outside the cochlea while the apical portion is inserted only to the region corresponding to the preserved residual hearing. This approach similarly preserves the option of deeper insertion should hearing loss progress (Lenarz et al., 2019).

Studies have shown that EAS significantly enhances speech recognition in complex listening environments compared to traditional CIs without acoustic components (Büchner et al., 2009; Incerti et al., 2013; Kiefer et al., 2005; Turner et al., 2004). However, combining electric and acoustic stimulation may introduce interactions that reduce the benefits of speech perception, a phenomenon known as electric-acoustic masking (Imsiecke et al., 2019). Electric-acoustic masking has been documented in various studies, including animal experiments on auditory nerve fiber activity (Miller et al., 2009; Tillein et al., 2015) and compound action potentials (McAnally et al., 1997; McAnally and Clark, 1994; Nourski et al., 2007; Stronks et al., 2012, 2010). Human studies using electrocochleography (ECochG), evoked compound action potentials (ECAPs; Imsiecke et al., 2020; Koka and Litvak, 2017; Krüger et al., 2020a, 2020b), and psychophysical masking (Imsiecke et al., 2019, 2018; Kipping et al., 2020; Krüger et al., 2017; Lin et al., 2011) have also explored these interactions.

Research suggests that electric-acoustic masking arises from peripheral processes such as direct electrical depolarization of spiral ganglion neurons (electroneural stimulation) and electrical stimulation of hair cells (electrophonic stimulation). Early studies primarily investigated electrophonic and acoustic stimulation interactions in animal models (Fráter, 2019; McAnally et al., 1997, 1993; McAnally and Clark, 1994; Stronks et al., 2013). Previous studies in EAS users suggested that electric stimulation configurated with long phase durations, amplitude modulation (Kiefer et al., 2001; Wilson et al., 1991), or rate coding (Nogueira et al., 2019; Wouters et al., 2015), could trigger electrophonic excitation. However, recent findings indicate that electric-acoustic masking can occur independently of pulse rate and phase duration (Kipping et al., 2020).

Psychoacoustic masking studies have explored how electric or acoustic hearing thresholds are elevated by either a simultaneous masker (Imsiecke et al., 2020, 2019; Kipping et al., 2020; Koka and Litvak, 2017; Krüger et al., 2017; Lin et al., 2011) or a preceding masker stimulus (Imsiecke et al., 2018), using pure tones for acoustic stimulation and unmodulated pulse trains for electric stimulation. Two main types of masking have been identified: electric masking of acoustic tones (“electric masking”) and acoustic masking of electric stimuli (“acoustic masking”). Threshold elevation (TE), defined as the difference between masked and unmasked thresholds, depends on acoustic frequency and electrode intra-cochlear location (Koka and Litvak, 2017; Lin et al., 2011). Krüger et al. (2017) found that electric-acoustic masking is influenced by the intra-cochlear distance between the sites of electric and acoustic excitation. Electric masking exhibits strong place dependence, affecting only acoustic tones located near the electrode contact based on cochlear tonotopic mapping. Rather than altering the acoustic tone itself or its transduction at the organ of Corti, this masking is thought to arise from a refractory state induced in auditory nerve fibers by the electrical stimulus, which transiently reduces their probability of responding to the subsequent or simultaneous acoustically evoked excitation (electroneural mechanism; Kipping et al., 2020; Krüger et al., 2017). In contrast, acoustic masking extends to more distant electrodes, allowing a single stimulating electrode to probe a broad range of the low-frequency hearing region simply by varying the acoustic masker frequency – an important advantage for diagnostic applications where electrode placement is inherently limited to a fixed location. These findings align with previous studies (Koka and Litvak, 2017; Lin et al., 2011) and have been replicated in subsequent research (Imsiecke et al., 2020, 2019; Kipping et al., 2020).

While these studies have established electric-acoustic interactions for electrical stimulation delivered via electrodes placed within the cochlea, it remains unknown whether comparable interactions arise when electrical stimulation is applied at or outside the cochlea. This study therefore examines both extra-and intra-cochlear electric-acoustic masking in individuals with CIs who use EAS, i.e., who receive electrical stimulation via the CI while retaining residual low-frequency acoustic hearing.

Accurate diagnosis of hearing loss is essential in determining a child’s residual hearing and selecting appropriate treatments like CIs. Behavioral audiometric techniques can estimate hearing thresholds but are not always feasible for children under 3.5 years of age, a critical period for brain development and optimal CI outcomes. Objective physiological measures, such as otoacoustic emissions (OAEs; Gong et al., 2020), electrocochleography (ECochG; Gibson, 2017; Koka and Litvak, 2017; Minaya and Atcherson, 2015), auditory brainstem responses (ABRs; Valderrama et al., 2014), auditory steady-state responses (ASSRs; Alaerts et al., 2010; Galambos et al., 1981; Gorga et al., 1988; Hayes and Jerger, 1982; Stapells et al., 1984), and cortical auditory evoked potentials (CAEPs; Van Dun et al., 2012) provide insight into auditory processing. However, assessing low-frequency hearing remains challenging due to noise susceptibility and the limitations of current diagnostic approaches, as diagnosing low-frequency hearing below 500 Hz, and even below 1 kHz, is challenging using existing methods that rely solely on either acoustic or electrical stimulation.

A promising approach for diagnosing residual hearing involves leveraging electric-acoustic interactions. Previous research suggests these interactions occur only when some degree of low-frequency acoustic hearing is preserved (Imsiecke et al., 2019). Consequently, evaluating electric-acoustic interactions could provide an objective means of assessing residual low-frequency hearing, which remains difficult to measure reliably because current clinical methods are particularly susceptible to noise at low frequencies. One potential application is using acoustic masking as an indicator of low-frequency hearing. Unlike acoustic masking, electric masking is spatially restricted to the cochlear region immediately adjacent to the stimulating electrode. Assessing the entire low-frequency hearing region using electric masking would therefore require multiple electrodes distributed across the apical portion of the cochlea. Such an approach is impractical, particularly for extra-cochlear stimulation, where electrode placement is inherently limited to a single fixed location, for example at the round window (RW). In contrast, acoustic masking allows the frequency of the acoustic masker to be varied while the electric probe is delivered through a single fixed electrode, thereby enabling assessment across a broad range of the low-frequency region.

However, previous studies (Imsiecke et al., 2019; Kipping et al., 2020; Krüger et al., 2017) have focused on interactions between low-frequency sounds and electric stimulation delivered via intra-cochlear electrodes. For a diagnostic device that aims to assess low-frequency auditory sensitivity without introducing additional trauma to the cochlea, electrical stimulation from an extra-cochlear location, such as the RW, would be preferable. In certain CI users, including those with partially inserted electrodes, an electrode may already be positioned at the RW, providing the opportunity to investigate electric-acoustic interactions originating from the basal cochlea or the RW region.

Gaining deeper insights into these interactions could lead to the development of innovative diagnostic and rehabilitation strategies. However, stimulation near the RW may inadvertently activate adjacent structures, such as the facial nerve (FN), the vestibular nerve (Bordure et al., 1989), middle ear muscles or other structures, leading to side effects (SEs; e.g., Bahmer et al., 2017). For example, facial nerve stimulation (FNS) occurs in approximately 5.6% of CI users (Van Horn et al., 2020). It is well-documented that electrical stimulation administered at the RW (e.g., Fourcin et al., 1983) or via electrodes placed externally around the cochlea (e.g., Banfai et al., 1984) can trigger auditory sensations. Furthermore, stimulation with electrodes near the RW has been employed for tinnitus suppression (e.g., Kheirkhah et al., 2024). However, additional research is required to ascertain whether specific configurations of electrical stimulation delivered to the RW or nearby locations in the basal region of the cochlea can evoke sound sensations without causing unintended SEs and whether these sensations can be masked by acoustic stimuli.

The flow of current into and out of the cochlea plays a central role in shaping auditory perception as well as potential SEs. To better understand these mechanisms, current distribution can be modeled, as demonstrated in previous work (Vanpoucke et al., 2004). Such models allow formulating and testing hypotheses on how current behavior in the cochlea contributes to acoustic masking. It is possible that acoustic stimulation may prevent electrical stimulation from activating low-frequency nerve fibers at the apex by inducing a refractory state, thereby increasing detection thresholds of electric stimulation in the presence of an acoustic masker.

This study investigates acoustic masking of electric stimuli through psychophysical experiments. The objectives are: (i) to characterize simultaneous ipsilateral electro-acoustic masking in EAS users under varying pulse rates, phase width, and stimulation types, and (ii) to examine auditory perception and SEs of electric stimulation in the basal region. Furthermore, this study examines current flow when activating electrodes near the RW compared to intra-cochlear electrodes. Trans-impedance measurements (TIMs), which evaluate the ratio between voltage and injected current within the cochlea, have been extensively used to study intra-cochlear current distribution, FN excitation patterns, and tissue growth (de Rijk et al., 2020; Saoji et al., 2024). These measurements in combination with a CI lumped model are particularly useful for assessing the spread of electrical stimulation and current flow in the cochlea (Vanpoucke et al., 2004). The results of the present study could facilitate the development of a novel diagnostic device for low-frequency acoustic hearing based on measured masking of electric stimulation delivered through extra-cochlear electric stimulation.

## 2. Methods

### 2.1. Subjects

Twelve EAS users with post-lingual deafness and residual ipsilateral acoustic hearing in the low frequency range participated in the study. Table 1 presents the patient’s demographic data as well as the respective experimental settings for each individual. At time of the study, participants age ranged between 24 and 83 years (mean 59.17) and had CI experience from 0.58 to 16.08 years (mean 10.25) in the implanted ear. All subjects participating in the study had Nucleus CIs with 22 electrodes. Seven subjects received a Nucleus CI24REH CI (Cochlear Ltd., Macquarie Park, NSW, Australia) with a 14.5 mm “Hybrid-L” electrode array. One additional subject was implanted with a Nucleus CI624 (Cochlear Ltd.) featuring a 19.1 mm “Slim 20” electrode array, which was only partially inserted as described by Lenarz et al. (2013), with the first four contacts located outside the cochlea and the remaining contacts inserted intracochlearly. Furthermore, three subjects received a Nucleus CI622 (Cochlear Ltd.) with a “Slim Straight” electrode array of up to 25 mm in length, and one subject was implanted with a Nucleus CI522 (Cochlear Ltd.) also featuring a 25 mm “Slim Straight” array.

**Table 1:**
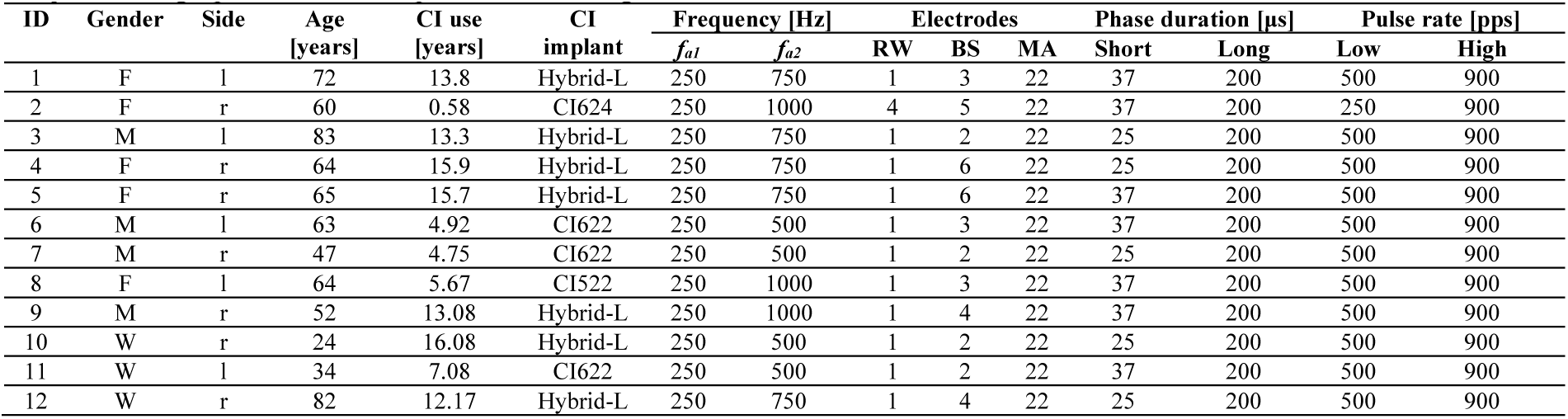
Subject demographic data and experimental configurations.

| ID | Gender | Side | Age<br>[years] | CI use<br>[years] | CI<br>implant | Frequency [Hz] |  | Electrodes |  |  | Phase duration [μs] |  | Pulse rate [pps] |  |
| --- | --- | --- | --- | --- | --- | --- | --- | --- | --- | --- | --- | --- | --- | --- |
| | | | | | | $f_{a1}$ | $f_{a2}$ | RW | BS | MA | Short | Long | Low | High |
| 1 | F | l | 72 | 13.8 | Hybrid-L | 250 | 750 | 1 | 3 | 22 | 37 | 200 | 500 | 900 |
| 2 | F | r | 60 | 0.58 | CI624 | 250 | 1000 | 4 | 5 | 22 | 37 | 200 | 250 | 900 |
| 3 | M | l | 83 | 13.3 | Hybrid-L | 250 | 750 | 1 | 2 | 22 | 25 | 200 | 500 | 900 |
| 4 | F | r | 64 | 15.9 | Hybrid-L | 250 | 750 | 1 | 6 | 22 | 25 | 200 | 500 | 900 |
| 5 | F | r | 65 | 15.7 | Hybrid-L | 250 | 750 | 1 | 6 | 22 | 37 | 200 | 500 | 900 |
| 6 | M | l | 63 | 4.92 | CI622 | 250 | 500 | 1 | 3 | 22 | 37 | 200 | 500 | 900 |
| 7 | M | r | 47 | 4.75 | CI622 | 250 | 500 | 1 | 2 | 22 | 25 | 200 | 500 | 900 |
| 8 | F | l | 64 | 5.67 | CI522 | 250 | 1000 | 1 | 3 | 22 | 37 | 200 | 500 | 900 |
| 9 | M | r | 52 | 13.08 | Hybrid-L | 250 | 1000 | 1 | 4 | 22 | 37 | 200 | 500 | 900 |
| 10 | W | r | 24 | 16.08 | Hybrid-L | 250 | 500 | 1 | 2 | 22 | 25 | 200 | 500 | 900 |
| 11 | W | l | 34 | 7.08 | CI622 | 250 | 500 | 1 | 2 | 22 | 37 | 200 | 500 | 900 |
| 12 | W | r | 82 | 12.17 | Hybrid-L | 250 | 750 | 1 | 4 | 22 | 25 | 200 | 500 | 900 |

The shorter Hybrid-L electrode array covers only the basal part (generally not more than 75% of the basal turn) of the cochlea to preserve low frequency hearing and it is specifically designed for CI candidates with residual hearing (Lenarz et al., 2013, 2009). Longer electrode arrays are implanted for patients with more severe hearing loss. With the discontinuation of the commercialization and manufacturing of the Hybrid-L electrode array and other short electrode arrays, the concept of partially inserting the electrode array in subjects with residual hearing became established (Lenarz et al., 2019). All subjects participating in the study had residual hearing.

Figure 1 presents the clinical audiograms of all subjects who participated in the study. Most subjects exhibited mild to moderately severe hearing loss up to 0.5 kHz (with the exception of subject ID 8) and profound hearing loss in the high-frequency range above 2 kHz (except for subject ID 3). In addition to residual hearing, subjects were required to have an electrode contact positioned near or directly on the RW to enable extra-cochlear stimulation. This placement was verified using cone beam computed tomography (CBCT) scans acquired during electrode insertion; for two subjects (IDs 01 and 04), no CBCT was available, and X-ray imaging was used instead. For one further subject (ID 10), CBCT was acquired but its image quality was insufficient to reliably visualize the RW niche, and a processed X-ray image was used instead. For these three subjects, RW electrode position was estimated based on the known geometry of the implanted electrode array (Hybrid-L, or Slim Straight; Cochlear Ltd.) together with the manufacturer-specified inter-electrode distances, following the protocol of Thormählen et al. (2024). An example of this is shown in Figure 2 for subject ID 11, who was implanted with a “Slim Straight” electrode array. Additional images of other subjects with different electrode types, imaging modalities, and localization approaches are provided in the appendix (Figure A 1).

**Figure 1.**
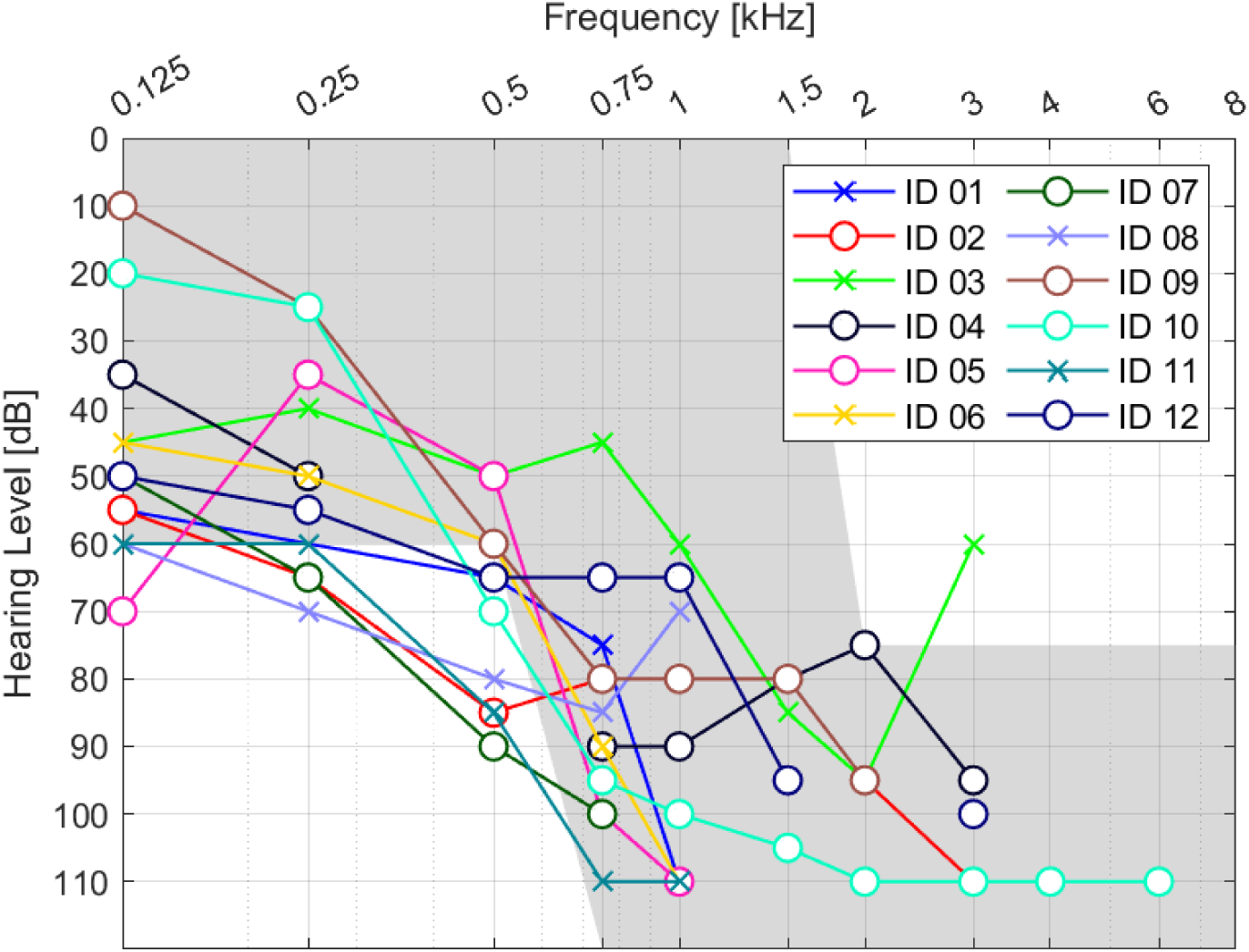
Clinical audiograms for each study participant. For each participant, the audiogram of the study ear is shown. The symbol indicates whether the study ear was the left (‘x’) or right (‘o’) ear. The grey shaded area represents the audiometric candidacy range for cochlear implant (CI) recipients using electric-acoustic stimulation (EAS).

**Figure 2.**
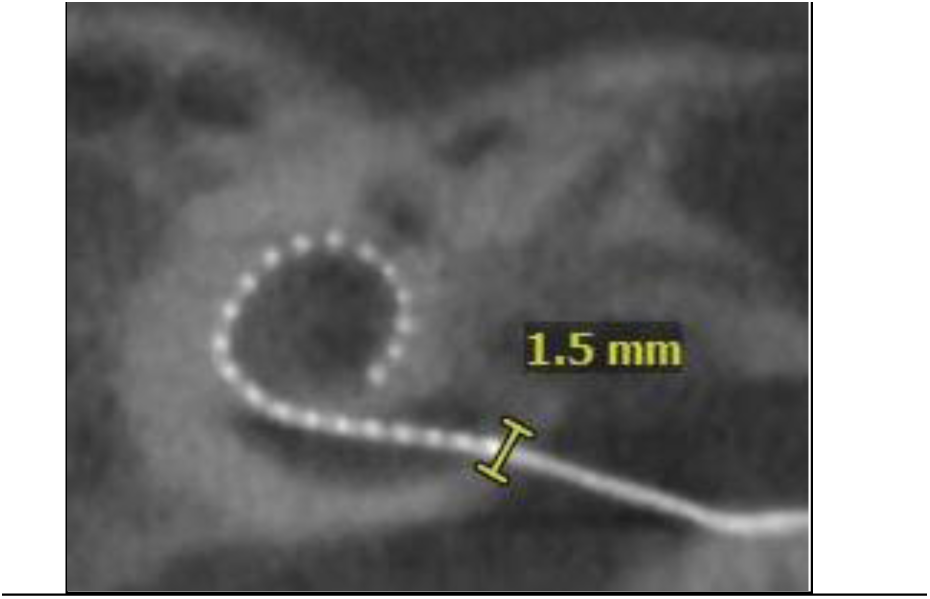
Illustration of a cone beam computed tomography (CBCT) image of subject ID 11. In this case, the electrode array is the Slim Straight (CI622). The yellow mark indicates the location of the RW, where the most basal electrode is visible. The RW’s diameter is approximately 1.5 mm.

All subjects provided written informed consent, as approved by the Institutional Review Board of the Hannover Medical School, Germany. Participation in the study was voluntary, and no financial compensation was offered.

### 2.2. Stimuli

Acoustic and electric stimulation was controlled with a custom made Matlab (The MathWorks, Inc., Natick, USA) script running on a desktop PC. A schematic representation of the experimental setup to conduct the experiments is presented in Figure 3.

**Figure 3.**
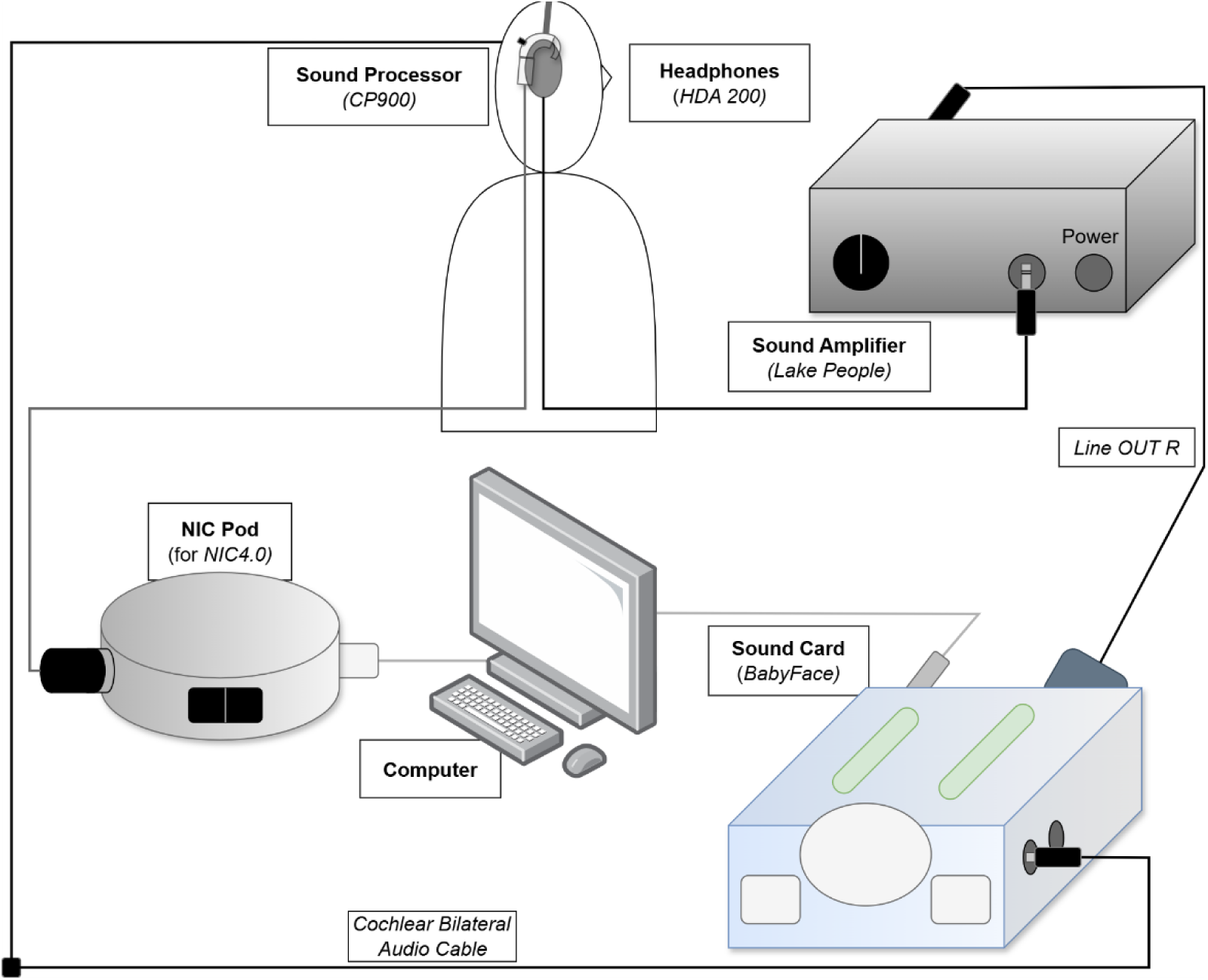
Diagram of the system architecture used for electrical-acoustic stimulation during the study. Acoustic signals are processed by a personal computer (PC) through an external RME Babyface sound card, amplified using a Phone-Amp G103 headphone amplifier, and delivered via HDA-200 headphones. Electrical signals are transmitted using a Cochlear CP900 processor, which is connected to the PC via a Cochlear Programming Pod.

#### 2.2.1. Acoustic stimuli

Acoustic stimuli consisted of ramped pure tones at several acoustic frequencies *fa* and broadband Gaussian noise spanning 20 Hz to 8 kHz. Pure tones were used to examine the frequency dependence of electric-acoustic masking, whereas Gaussian noise provided simultaneous stimulation across the clinically relevant frequency range. The broadband stimulus was included to maximize masking by engaging a wider bandwidth of the auditory system than could be achieved with frequency-specific pure-tone stimulation. The acoustic stimuli are used as masking signals with a duration of 500 ms, including 50 ms of on-and off-ramps. The maskers were always presented at the most comfortable level (MCL). A schematic representation of the acoustic signal is shown in Figure 4 (b). Acoustic signals were processed with an external RME Babyface sound card (Audio AG, Hainhausen, Germany), amplified by a Phone-Amp G103 headphone-amplifier (Lake People electronics GmbH, Konstanz, Germany) and delivered via HDA-200 headphone (Sennheiser electronic GmbH & Co. KG, Wedemark, Germany). Figure 3 shows a visualization of the setup. Calibration was carried out with a Brüel & Kjær Sound Level Meter Type 2250 and an Artificial Ear Type 4153 (Brüel & Kjær Vibro A/S, Nærum Denmark). For each subject, a pure tone with acoustic frequency *fa1* = 250 Hz and a Gaussian noise (*GN*) were used for the experiment. A second pure tone with a frequency *fa2* of either 500 Hz, 750 Hz or 1000 Hz depending on the participant’s residual acoustic hearing was used for the experiment. The chosen acoustic frequencies for each subject are listed in Table 1.

**Figure 4.**
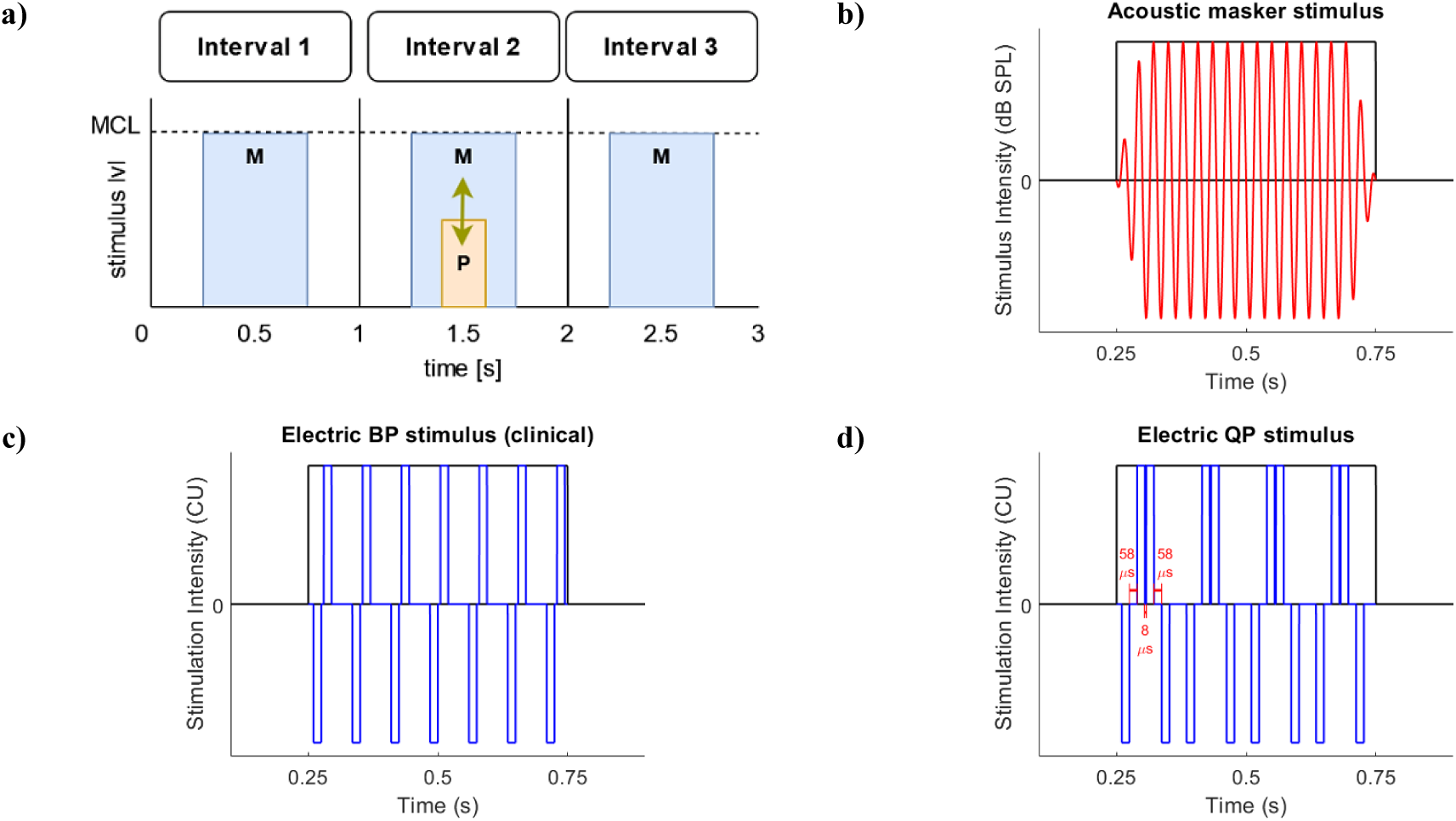
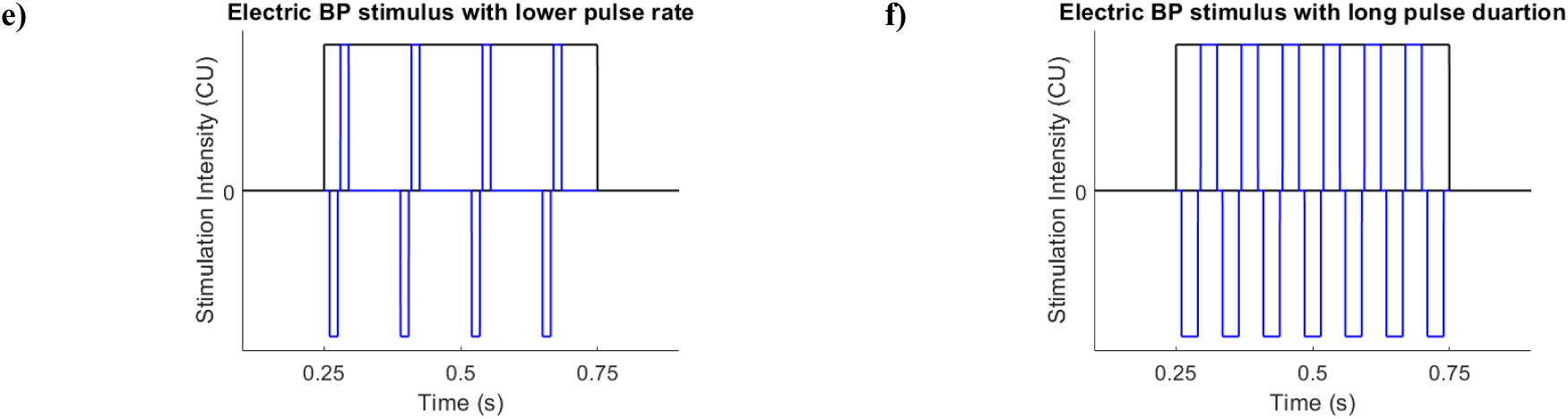
Stimuli used in the three-interval forced-choice masking paradigm. **(a)** Schematic of a single trial consisting of three 1-second intervals. Masker stimuli (M) were presented at the most comfortable loudness level (MCL) in all intervals, while the electric probe stimulus (P) was adaptively varied and randomly presented in one of the intervals. **(b)** Acoustic masker stimuli consisted of 500 ms ramped pure tones or Gaussian noise, including low-frequency (250 Hz), high-frequency (individualized, typically ∼750 Hz), and white-band noise (20 Hz – 8 kHz). **(c)** Electric biphasic (BP) stimulation under clinical settings (pulse rate: 900 pps; phase width: 25 or 37 µs, subject-dependent). **(d)** Electric quadraphasic (QP) stimulation using the same parameters as in (c), applied in a pseudo-triphasic (TP) configuration. **(e)** Electric BP stimulation with reduced pulse rate (500 pps; 250 pps for subject ID 2) and clinical phase width. **(f)** Electric BP stimulation with extended phase width (200 µs) and clinical pulse rate. All electrical probe stimuli were unmodulated pulse trains of 200 ms duration. The inter-phase gap (IPG) was set to 8 µs for all configurations, except for QP stimuli (58 µs).

Some participants (ID 1, 8 and 10) had residual hearing and CIs on both sides. For these individuals, the side with the better performance was chosen for the experiment. The difference between the acoustic stimulus presentation level on the test side and the hearing threshold on the opposite side was always less than 40 dB, within the known attenuation range for bone-conduction cross-hearing (Stenfelt, 2012). All participants reported no auditory perception on the side opposite to the one being tested.

#### 2.2.2 Electric stimuli

The probe signals lasted 200 ms and were presented at different intensities using a three-interval forced-choice procedure. The electrical stimuli were unmodulated pulse trains of charge-balanced biphasic (BP) or quadraphasic (QP) pulses, with interphase gaps (IPGs) of 8 and 58 μs, respectively. Two BP pulses were used to create a QP pulse that mimics a pseudo-triphasic (TP) pulse (Carlyon et al., 2013; Macherey et al., 2010). The NIC4 Cochlear Research Platform (Cochlear Ltd., Macquarie Park, NSW, Australia) only supports symmetrical pulses with opposite polarity. When QP pulses are used, these two BP pulses are separated by a minimum phase difference of 8 μs. Due to technical limitations of the platform, the IPG for QP pulses was set to 58 µs (see Figure 4d). All pulses had a cathodic-leading phase, which has been shown to be effective for electric stimulation in previous studies (Bahmer et al., 2017). Electric stimulation was delivered via a Cochlear CP900 processor connected to a Cochlear Programming Pod. Figure 3 shows a visualization of the setup. Before testing, the correct stimulation was verified using a Cochlear DIET box and a PicoScope 5443A oscilloscope (Pico Technology Ltd., Cambridgeshire, UK) connected to a cochlear reference implant. Electric stimulus levels were defined in current units (CU) which are related to current levels in μA as described in equation (1). The maximum stimulation level was 255 CU, corresponding to 1.75 mA.

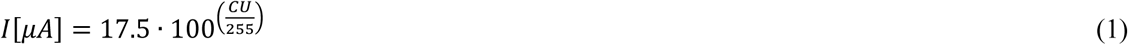

For testing, the pulse rate was set to either 250, 500 or 900 pulses per second (pps). The pulse rates of 250 and 500 pps were classified as “short”, whereas 900 pps was classified as “long”. Note that the selected 900 pps coincides with the clinical setting of the participants. The duration of the phases was labelled as “short” which corresponded with 25 μs or 37 μs mimicking the clinical setting. For the “long” configuration the phase width was 200 μs. This value does not reflect typical clinical CI fitting parameters, which commonly use considerably shorter phase durations, but was instead chosen as an experimental condition to investigate the effect of long phase durations on electric-acoustic masking. Long phase durations require lower current amplitudes to reach equivalent charge levels and result in longer per-pulse stimulation, and have been used in previous studies investigating their influence on electric-acoustic interaction, where phase durations of 200 or 400 μs (depending on the subject) were similarly tested as “long” conditions (Kipping et al., 2020). A schematic representation of the different electrical signals is given in Figure 4 (c) - (f).

Electrical stimuli were delivered via the electrode placed directly on or at the RW, an active basal electrode that was active in the clinical setting of the subject (BA), and via the most apical electrode (AP), which was electrode 22 in all study participants. The BA electrode is selected so that it is the first basally active electrode, unless this is the RW electrode. In this case, the BA electrode would be the next one.

Additionally, the impedances of individual electrodes were measured to ensure the stimulation was not “out-of-compliance”. Stimulation was “out-of-compliance” if the stimulation current exceeded

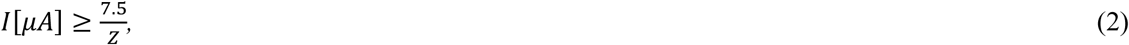

where Z is the measured impedance value in Ω.

### 2.3. Procedure

An experimental session included multiple pre-measurements followed by the main experiment. The session lasted four to five hours, including testing time and additional breaks. During the pre-measurements, the MCLs for both electric and acoustic stimuli were identified. Furthermore, a loudness-balancing procedure was performed to ensure the MCLs of the acoustic stimuli were perceived as equally loud.

#### 2.3.1. Audiogram

To determine the residual hearing threshold on the measurement day, participants completed an audiogram in a dedicated acoustic chamber designed for clinical audiometric testing and an audiometer (Audio-Ton, Hamburg, Germany). HDA-300 headphones (Sennheiser electronic GmbH & Co. KG, Wedemark, Germany) were used. A response button was used to indicate a perceived tone by the study participants.

Testing began with the best acoustic hearing side (the ear selected for the study) and was followed by the contralateral ear to check for cross-hearing effects (see 2.2.1. Acoustic stimuli). The procedure started with a 1000 Hz tone. Frequencies were then progressively lowered to 125 Hz following the standard audiometric frequencies and subsequently increased back to 1000 Hz, continuing to the maximum tested frequency of 8000 Hz. When a tone was perceived and reported via the response button, the same tone was reduced by 20 dB and then gradually increased again to confirm the previously determined threshold.

#### 2.3.2. Trans-impedance measurements

Trans-impedance measurements (TIMs) were performed to investigate current flow when stimulating electrodes located around the RW, both inside and outside the cochlea. Comparisons between TIMs were made when stimulation was delivered through extra-and intra-cochlear electrodes.

Figure 5 above illustrates a TIM obtained when the RW electrode was stimulated. In this context, impedance was defined as the ratio of the recorded voltage at a given electrode to the current injected through the stimulating electrode.

**Figure 5.**
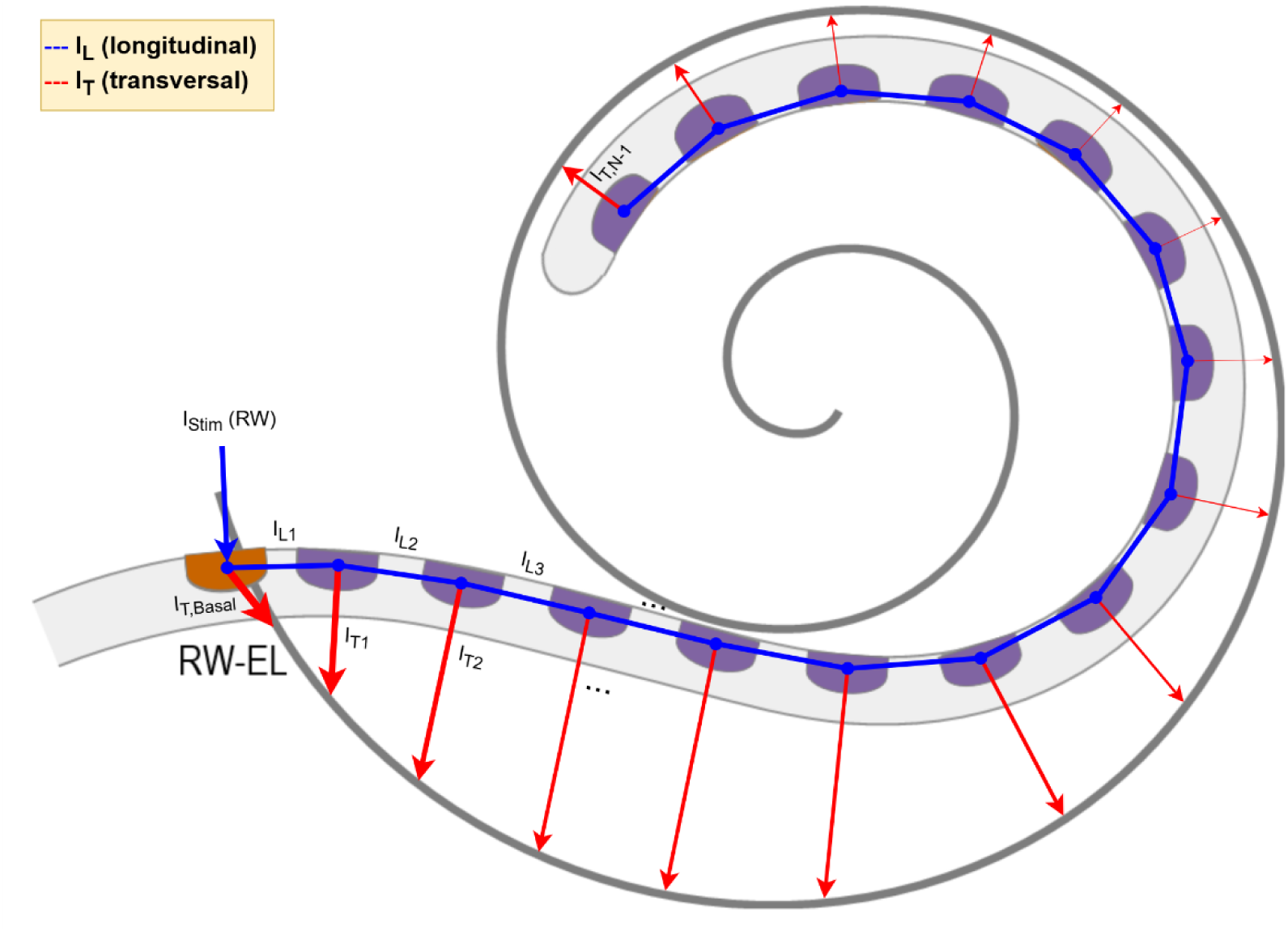
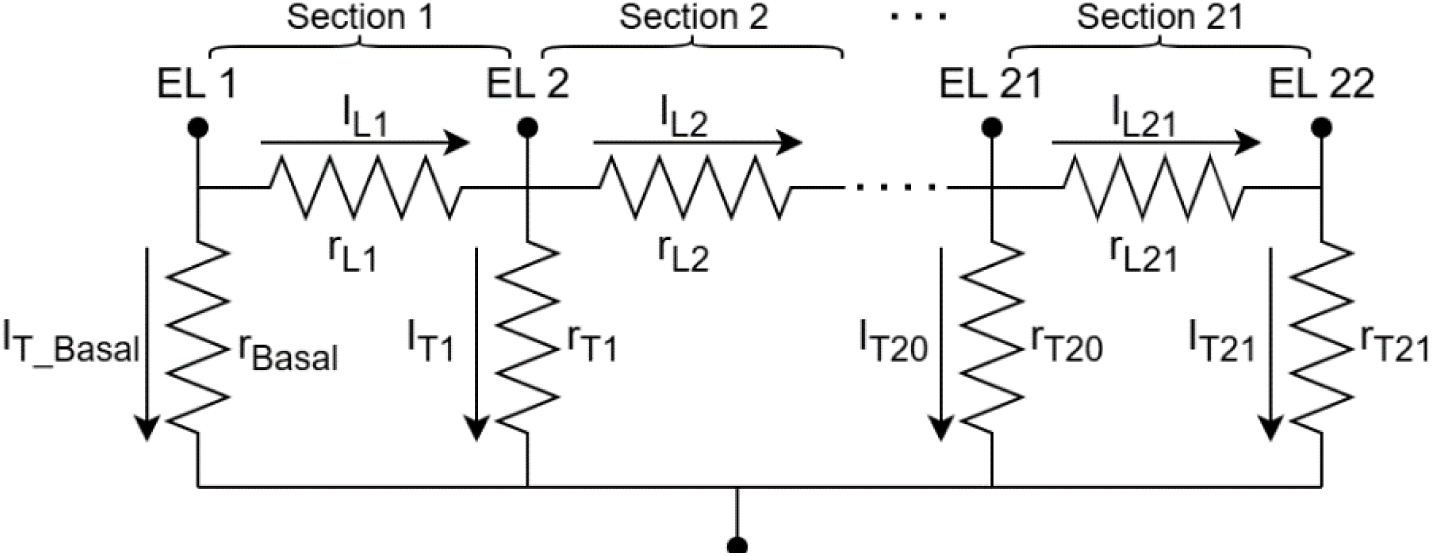
**Above**: Schematic illustration of a trans-impedance measurement (TIM) obtained when the round window electrode (RW-EL, orange) is stimulated. Stimulation is applied to the RW-EL and voltage is recorded at the other electrode contacts (purple), from which impedance is derived as the ratio of recorded voltage to injected current. The schematic additionally illustrates the underlying current flow: current propagates longitudinally (*I_L_*) through the scala tympani along consecutive intra-cochlear contacts, while proportion of current leaks transversally (*I_T_*) at each contact toward the cochlear wall and extra-cochlear space. Arrow length reflects the pattern of transversal current predicted by the lumped-parameter model during RW stimulation: large near the stimulation site (*I_T_*_,*Basal*_), decreasing toward mid-cochlear contacts, and increasing again toward the most apical contacts (see also 4.1. Trans-impedance measurements and simulated cochlear current flow). Note that this sketch is schematic and depicts 16 electrode contacts for illustrative purposes; the electrode array used in the present study (Cochlear Ltd.) comprised 22 contacts. **Below**. Electric current flow model adapted from Vanpoucke et al. (2004). The propagation of current through the cochlear tissue is modeled by a leaky transmission line network. The network consists of 21 sections, each of which contains transversal and longitudinal resistances. An additional basal resistance completes the network.

TIMs were acquired using the CustomSound EP 6 software (CS6; Cochlear Ltd., Macquarie Park, NSW, Australia). Analyses focused on the slope of the TIMs, comparing stimulations at the RW electrode and at the intra-cochlear electrode EL12. The slope decay was estimated from the first two electrodes adjacent to the stimulating electrode, in order to highlight local effects. For RW stimulation, this involved the cochlear electrodes directly neighboring the RW electrode, whereas for EL12 stimulation, electrodes 11–10 (towards the base) and 13–14 (towards the apex) were considered.

Stimulation currents were set to 109 CU as specified by CS6, except for subject ID 2, where 192 CU was applied. To enable comparability, TIMs from ID 2 were scaled accordingly.

#### 2.3.3. Threshold-comfort level of electric and acoustic stimuli and loudness balancing

Threshold and comfort levels were measured for each subject when an electrode located close to the RW was stimulated. Additionally, SEs were tracked when increasing the current at this particular electrode. Threshold and comfort levels were also measured with intra-cochlear electrodes (BA and AP) for comparison and with acoustic stimulation.

Subjects rated perceived loudness on a scale from 1 to 10, corresponding to “extremely quiet” to “extremely loud.” Stimulation began at 0 CU for electric stimuli and at 0 dB SPL for acoustic stimuli. The level was gradually increased to determine the threshold level (T-Level; corresponding to the transition from level 0 to 1), the MCL (level 6), and the uncomfortable loudness level (UCL; transition from level 7 to 8). After reaching the UCL, subjects were asked to self-adjust the electrical and acoustic MCL using a rotary encoder (Griffin PowerMate, Griffin Technology, Irvine, USA). The MCL was estimated as the average of the self-adjusted and scaled comfort levels. Because the MCL was derived from two separate loudness judgments, one obtained during the initial ascending scaling procedure and one obtained via subsequent self-adjustment, whereas the UCL was determined from a single ascending judgment, the resulting MCL and UCL values could be numerically close. In some individual cases, they could even be nearly identical, reflecting the inherent variability of subjective loudness judgments near the upper end of the dynamic range.

Participants were also asked to report any adverse SEs, particularly in response to electric stimulation at the RW electrode. To support this, they completed a standardized form specifying the location at different parts of the head or the body, the type, and the intensity of any perceived effect. The intensity of the SE was recorded on a scale of 1 (barely noticeable) to 5 (intolerable).

#### 2.3.4. Loudness balancing of acoustic stimuli

Subsequently, the approximately determined acoustic MCLs for the pure tones *fa₁* (low frequency at 250 Hz) and *fa₂* (subject-specific higher frequency; see Table 1), were pairwise loudness-balanced, as well as for the Gaussian noise. The procedure followed the loudness balancing method described by Krüger et al. (2017) and Kipping et al. (2020). A reference and an adjustable acoustic stimulus were alternately presented, and participants adjusted the latter to match the perceived loudness of the reference. Each pair was tested twice, starting from −10% and +10% of the rough MCL estimate for the adjustable stimulus. For the three acoustic conditions, adjacent frequencies were compared using the lowest frequency *fa1* as the initial reference.

#### 2.3.5. Masking experiment

First electric hearing thresholds of participants were measured using an adaptive three-interval forced-choice (3-AFC) procedure, as described in Lin et al. (2011), Krüger et al. (2017), and Kipping et al. (2020). Thresholds were measured under both masked and unmasked conditions. Each threshold was determined through a sequence of typically 30 to 60 trials.

A run involved the sequential presentation of three 1-second intervals, with the corresponding buttons highlighted on a graphical user interface. One of the intervals randomly contained the 200 ms electric probe stimulus, which the subject was asked to detect. In the masked configuration, additional 500 ms acoustic maskers were introduced in each interval. The acoustic masking experiment assessed the impact of the acoustic maskers on electric probes. The maskers were always presented at MCL, determined in the previous step, and all stimuli were temporally centered within their intervals. A graphical illustration of the 3-AFC procedure is shown in Figure 4 (a).

Within each run, the probe level was adaptively adjusted based on the subject’s responses using a one-up, two-down rule (Levitt, 1971). The initial stimulus level for the adaptive procedure was set to 8 CU and reduced by half after the first and second reversals. The procedure continued until four additional reversals were completed at the minimum step size of 2 CU. The probe threshold was then estimated as the mean of these final four reversals (Krüger et al., 2017). This procedure is expected to converge at the 70.7% correct response level on the psychometric function.

The experiment was conducted with various combinations of masker and probe stimuli, presented in randomized order. Since the analysis was particularly sensitive to unmasked thresholds, all unmasked conditions were tested twice, and the final threshold was calculated as the average of both runs. Test-retest reliability for unmasked electric probes showed a standard deviation of 2.12 CU. Due to time constraints, masked conditions were typically tested only once. Any run in which the standard deviation of the final four reversals exceeded 4 CU was excluded and repeated.

### 2.4. Analysis

#### 2.4.1. Dynamic ranges

Threshold elevation (TE) caused by simultaneous masker presentation was calculated as the difference between the masked threshold *θₘ* and the unmasked threshold *θᵤ*. To enable comparisons across subjects with varying degrees of hearing loss and different sensitivity to electrical stimulation, results were expressed relative to each subject’s dynamic range (DR) for the corresponding probe stimulus:

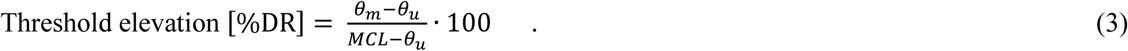

Figure 11 shows the DRs for electrical probes with a 200 ms duration.

#### 2.4.2. Simulated current flow model

A lumped-parameter model based on Vanpoucke et al. (2004) was used to simulate current flow from the stimulating electrodes using the measured TIM values (see Section 2.3.2. Trans-impedance measurements). The model has four layers representing distinct components of the current path: (1) output lines of the internal current sources, (2) the material of the electrode contacts, (3) the perilymph medium, and (4) the tissue of the extra-cochlear reference electrode.

Layer 3 implements a leaky transmission-line network to model current spread through cochlear tissues. The model contains sections, each comprising a transversal resistor (modeling the current pathway from the scala tympani (ST) back to the reference electrode through surrounding lateral wall and modiolar bone structures) and a longitudinal resistor (modeling the flow of current along the ST). The model provides relative current distributions between adjacent electrodes, expressed as transversal currents *IT* (leaving the ST) and longitudinal currents *IL* (within the cochlea, directed apically or basally).

For the present work, the model was adapted to an electrode array with *N* = 22 contacts (Figure 5 below). The adapted model is represented as an *N* – 1 chain of sections, each corresponding to the section between two successive contacts, with segment spacing determined by the implanted array. Each segment contains one node for the ST potential, connected by longitudinal resistances modeling current flow along the scala and transversal resistances to ground modeling current leakage toward the reference electrode. An additional basal resistance accounts for conductivity between the cochlear base and the reference electrode through extra-cochlear pathways (internal auditory canal, vestibular system, cochlear and vestibular aqueducts, and middle ear). Since these structures cannot be directly measured, this basal resistance is modeled as a single lumped element, and the corresponding transversal current (*IT*,*Basal*) represents current leaving the modeled network through this basal shunt rather than through a specific point along the ST – regardless of whether the stimulating electrode is located at the RW or at an intra-cochlear position (BA, AP).

Kirchhoff’s current law requires that, at every node, the current entering must equal the current leaving. Applied to the model, the current injected at the stimulating electrode (*IStim*) divides at each subsequent node into a fraction that leaves the ST transversally (*IT*) and a fraction that continues to propagate longitudinally (*IL*) — with the latter splitting into apically and basally directed components when the stimulating electrode is located at an intra-cochlear position. For the case illustrated in Figure 5 (above), where current is injected at the RW-EL, i.e., at the basal-most position of the modeled chain, all longitudinal current propagates in the apical direction only (see also Figure 5, above, for the notation):

- At the stimulating node:

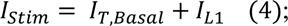

the injected current splits into what leaves through the basal shunt (*IT*,*Basal*) and what continues apically into the array (*IL*1).
- At each following node *k*:

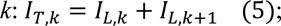

the transversal current leaving at that node equals the difference between what arrived from the more basal side (*IL*,*k*) and what continues onward (*IL*,*k*+1). For example, *IT*2 = *IL*2 − *IL*3.
- At the most apical node, which has no further node to propagate to, all remaining current must leave transversally:

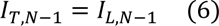

The same principle applies symmetrically for stimulation at an intra-cochlear electrode: current entering the node splits into a transversal component and two longitudinal components propagating in opposite directions (apically and basally), each obeying the same node-by-node current balance along its respective direction.

Because every unit of injected current eventually leaves the ST transversally at some position along the array – whether close to the stimulation site or, at the latest, at the terminal node in either direction – the transversal currents across all modeled positions sum to the total injected current (∑ *IT*,*k* = *IStim*, i.e., 100% when normalized). The longitudinal currents *IL*,*k*, in contrast, do not sum to a fixed value: they simply decrease step by step in each direction away from the stimulation site as current is progressively lost transversally, until reaching zero beyond the respective terminal node.

#### 2.4.3. Relation between current flow and masking

Acoustic masking occurs through interaction at the apex of the cochlea and may prevent electrical stimulation from exciting low-frequency nerve fibers at the apex by inducing a refractory state, thereby increasing TE in the presence of an acoustic masker. This was tested by correlating 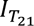 (current flow towards the apex) with TE for all acoustic maskers (*fa1, fa2, GN*), using both the BP stimulation (all subjects) and the optimized configuration (subjects perceiving T, M, and U levels without SE; see 2.3.3. Threshold-comfort level of electric and acoustic stimuli and loudness balancing).

#### 2.4.4. Relation between outflux current flow and side-effects

Outflux current flow toward the RW, and possibly extending beyond the cochlear boundaries towards the extra-cochlear space (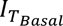), may be related to the perception and severity of SEs. For the partially inserted patient ID 2, this outflux current corresponds to the longitudinal current 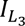 at EL4 (RW).

#### 2.4.5. Relation between residual hearing and masking

Residual hearing was additionally considered as a predictor of masking. Multiple linear regression (MLR) was performed using pure-tone (PT) or averaged (PTA) audiometric thresholds alongside model-derived current values and TE. For individual tones, PT values were taken from the audiogram; for white noise, a PTA from 125 Hz to 2000 Hz was used, corresponding to the maximum range perceived by subjects ID 2, 3, 4, 9, 10. Integrated masking data (mean from *fa1* up to *fa2* tones) was also computed for this PTA range.

All correlations and MLR were conducted for both absolute (CU) and relative (%DR) TE values.

## 3. Results

### 3.1. Trans-impedances and simulated cochlea current flow

Figure 6 illustrates the measured trans-impedances (in [Ω]) for each subject. Figure 6 (a) displays the trans-impedances when stimulating the CI electrode closest to the RW. A clear decrease in trans-impedance is observed in the apical direction with increasing distance from the RW. The measured slopes vary across subjects, depending on implant type and insertion depth, ranging from a minimum of 16.5 Ω/mm (ID 11) to a maximum of 2268 Ω/mm (ID 5). Note that electrode spacing varies slightly across subjects due to differences in array design, For comparison, Figure 6 (b) shows the trans-impedances when an intra-cochlear electrode near the center of the array (specifically, EL12) is stimulated. Here, trans-impedance also decreases in both apical and basal directions, but with a noticeably shallower slope than with RW stimulation, ranging from 108.3 Ω/mm (ID 8) to 1384.6 Ω/mm (ID 5). Notably, in subject ID 2, who has a partially inserted CI with EL 4 located at the RW, a distinct drop in trans-impedance is observed for electrodes basal to EL 6, suggesting current flow toward the extra-cochlear space.

**Figure 6.**
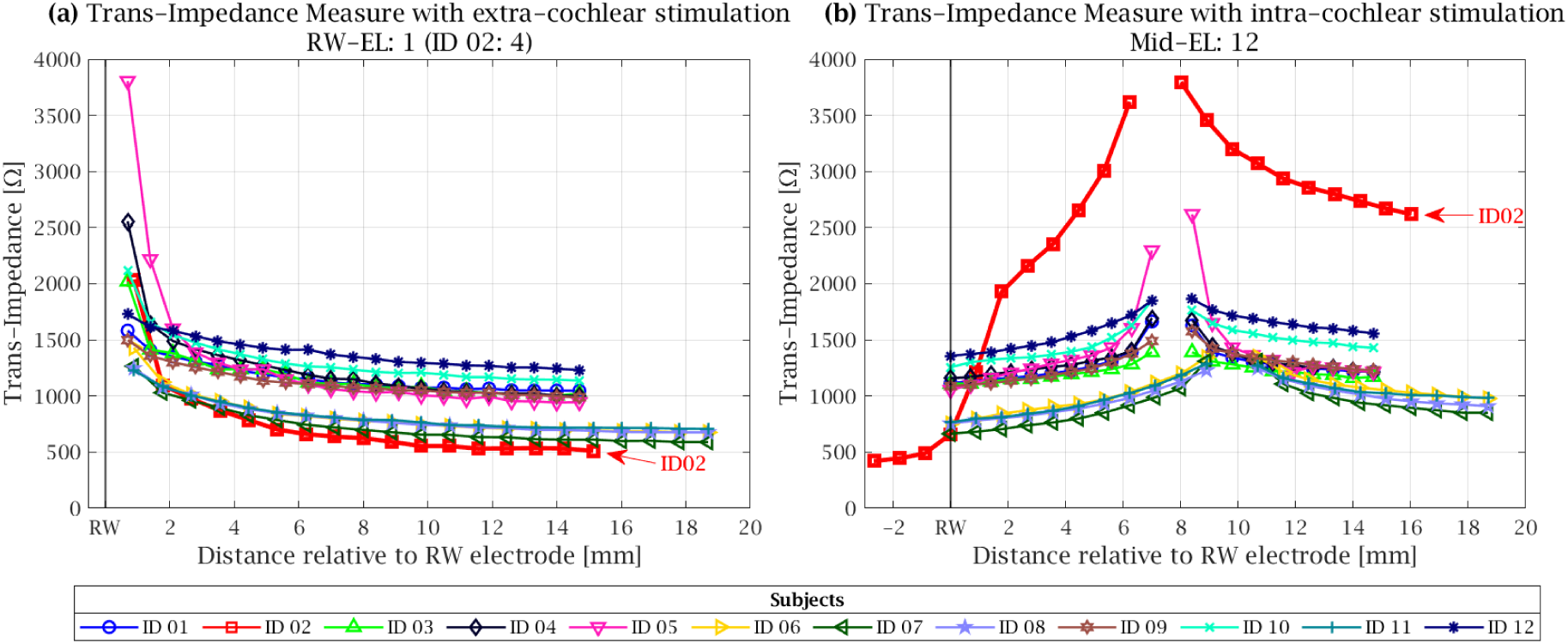
Trans-impedances in all subjects. **(a)** Stimulation at the most basal electrode (RW-EL1; ID 02: EL4). **(b)** Stimulation at a mid-array electrode (EL12). Each subject is represented by a unique combination of color and marker symbol, which is used consistently for all trans-impedance measurements across electrode locations. The x-axis shows the distance of each electrode contact from the round window (RW), which varies with the implanted array (Hybrid-L: 0.7mm, others: 0.89mm; see Table 1). Trans-impedances are highest near the stimulating electrode and decrease with distance. Measurements at the stimulating electrode were excluded, as they are dominated by the electrode impedance and do not reflect intra-cochlear current flow. All measurements were performed with a stimulation current of 109 CU (automatically set by CS6), except for subject ID 02, where 192 CU was used (automatically set by CS6). Trans-impedances of the partially inserted patient (ID 02) are indicated by separate arrows.

The model-based analysis of current flow (Figure 7) provides insight into stimulation patterns at different sites in and around the cochlea. It illustrates the modeled transversal (*IT* in % of injected current, reflecting current leaving the ST at a given axial position) and longitudinal (*IL* in %, reflecting current remaining inside the ST between adjacent positions) branch currents for stimulation with the RW, BA and AP electrodes (see 2.4.2. Simulated current flow model for the formal derivation).

**Figure 7.**
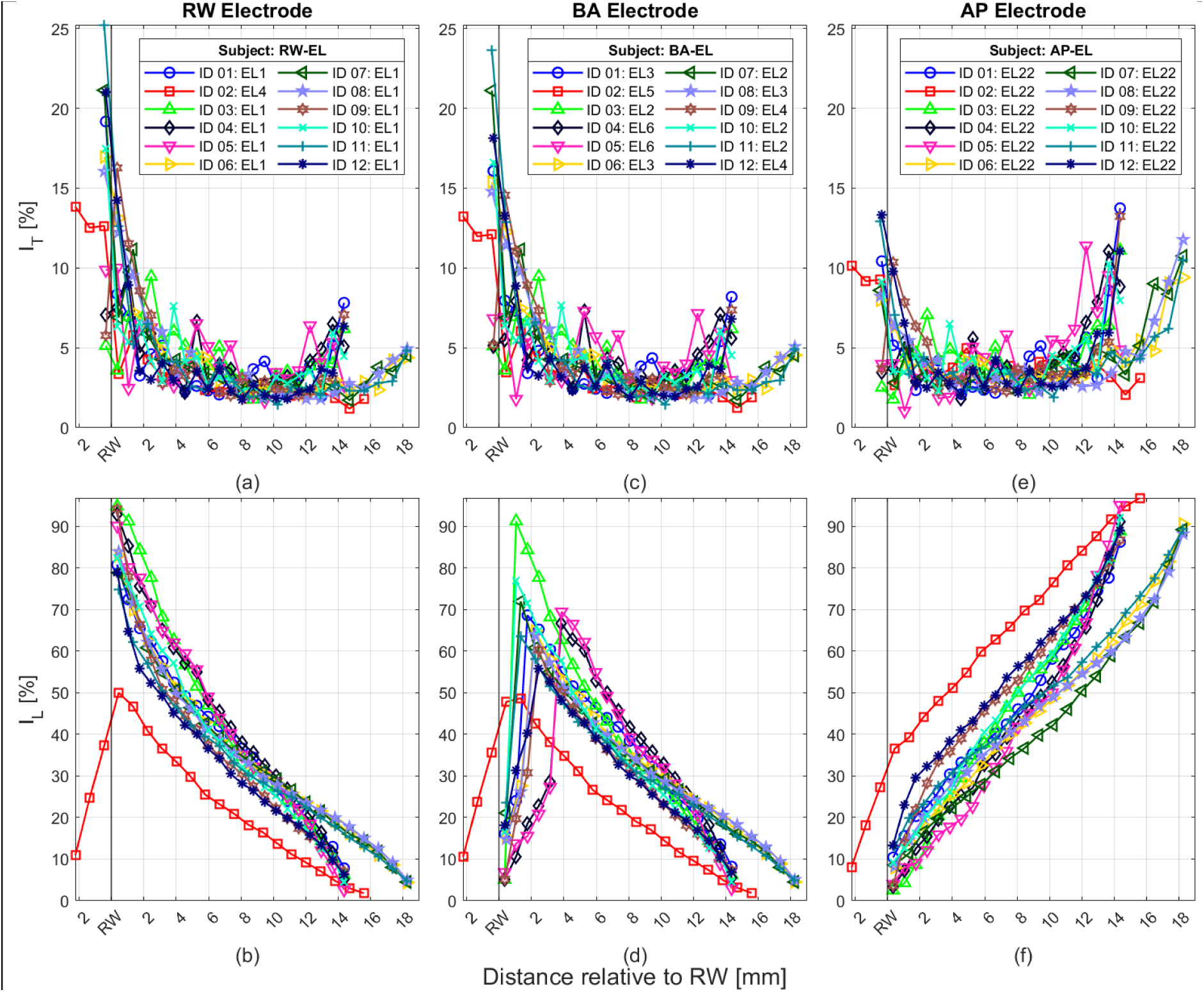
Transversal (*I_T_*) and longitudinal (*I_L_*) currents estimated across cochlear locations in (%) when electrical stimulation is delivered at different sites (RW: round window; BA: base of the cochlea; AP: apex of the cochlea). Panels (**a, c, e**) show transversal currents, and panels (**b, d, f**) show longitudinal currents for stimulation at the RW, the next basal active (BA), and the most apical (AP) electrode. Each subject is represented by a unique combination of color and marker symbol, which is used consistently for all data points across the cochlear electrode locations. Current values represent the percentage of transversal or longitudinal flow between electrode segments. According to the model, transversal currents represent current leaving the scala tympani (ST) toward the extra-cochlear space at each axial position, while longitudinal currents represent current propagating along the cochlear axis within ST. The RW electrode serves as the spatial reference, and inter-electrode distances vary with the implanted electrode array (see Table 1).

The transversal currents (Figure 7 (a), (c), (e)) present similarities in terms of its decay pattern and magnitude across subjects from the stimulating electrode to the adjacent ones in the basal region. Note that 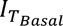 which is related to the current exiting the cochlea or outflux current is not directly related to a specific location in or outside the cochlea. Therefore, for all subjects except ID 2, the most basal transversal current (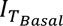; see also Figure 5 below) has been plotted at the location RW+Δ mm, where Δ is the inter-electrode spacing for each individual, to facilitate visualization. In subject ID 2, who had a partially inserted electrode array, 3 transversal currents are displayed at locations RW-Δ mm, RW-2Δ mm and RW-3Δ mm also to facilitate visualization. It is noteworthy that the largest transversal currents during RW and BA stimulation are observed at 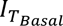 indicating a pronounced outflux current. Similarly, AP stimulation also results in substantial 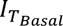 currents, again suggesting significant outflux. In the partially inserted CI subject ID 2, outflux current can be detected through transversal currents estimated from the extra-cochlear electrodes. These estimates indicate that, upon stimulation of the RW, BA, and AP electrodes (Figure 7), current flows outward from the cochlea.

For longitudinal currents (Figure 7 (b), (d), (f)), current flow patterns are also generally consistent across subjects. In most cases, stimulation at the RW electrode results in currents entering the cochlea, as demonstrated by large 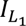 ; percentage currents. The partially inserted subject ID 2 again represents a notable exception. Stimulation provided at the RW electrode leads to a considerable outward current (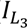 of up to 37%) and a comparatively lower inward current (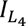 50% at RW+1). This inward current continues toward the more apical electrodes and decreases progressively, reaching almost 0% at the AP electrode. Similarly, stimulation at the BA electrode (EL5) in ID 2 produces a higher proportion of outward current (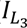), whereas in the other subjects, BS stimulation primarily generates branching longitudinal currents that demonstrate current flow towards the inside of the cochlea, characterized by a steep decline toward the RW electrode and a more gradual reduction in the apical direction.

Figure 8 presents the modeled current flow across all subjects when stimulation was delivered to the RW, BA and AP electrodes. Figure 8 (a) shows the proportion of current leaving the cochlea towards the extra-cochlear space (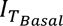). Current outflux was highest for RW electrode stimulation (median = 17.21%), followed by BA stimulation (median = 15.73%) and AP stimulation (median = 8.41%). A Friedman test revealed a significant effect of stimulation site (*χ*^2^(2) = 24.0, *p* < 0.001). Post-hoc Wilcoxon signed-rank tests with Holm correction confirmed significant differences between all electrode pairs (RW vs. BA, RW vs. AP, and BA vs. AP; all adjusted *p* = 0.0015), indicating a progressive decrease in extra-cochlear current flow from RW to BA to AP stimulation.

**Figure 8.**
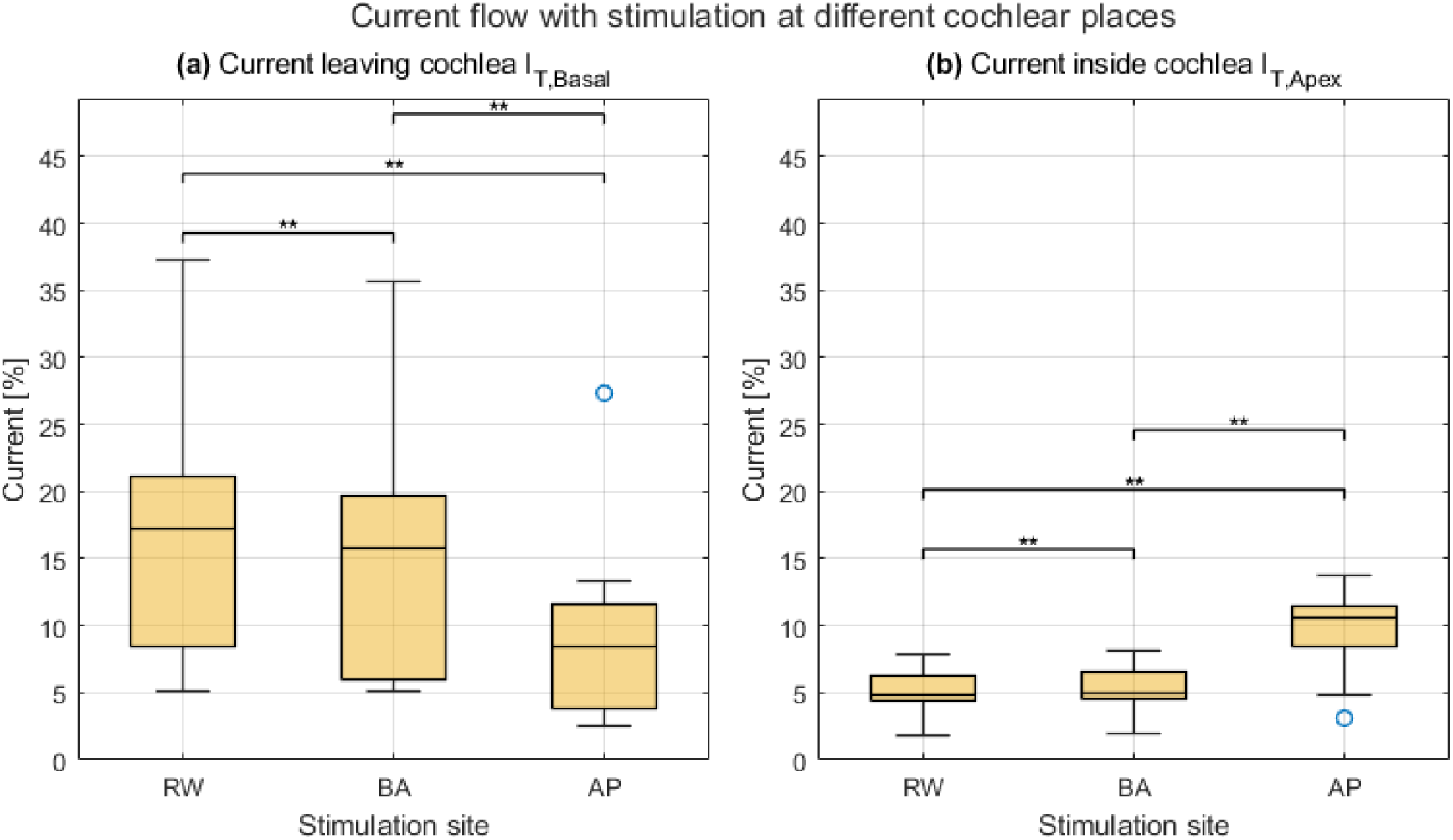
Box plots of modeled current flow during stimulation at different electrode sites: round window (RW), base (BA), and apex (AP), shown for all subjects. **(a)** Estimated current outflux, corresponding to the transversal basal current (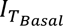) (see also Figure 5), except for subject ID 2 with a partially inserted CI, for whom the longitudinal current (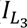) was used. **(b)** Estimated current flux at the apex of the cochlea (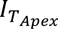). The currents represent relative current flows derived from the model shown in Figure 7. A Friedman test revealed significant differences between stimulation sites for both current measures (***p*** < 0.001). Post-hoc Wilcoxon signed-rank tests with Holm correction showed significant pairwise differences between RW, BA, and AP for both 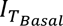 and 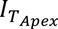 (all adjusted ***p*** = 0.0015).

In contrast, Figure 8 (b) illustrates the modeled current flowing towards the cochlear apex (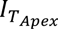). Stimulation at the AP electrode resulted in the highest intra-cochlear current flow towards the apex (median = 10.63%), compared with BA stimulation (median = 5.00%) and RW stimulation (median = 4.80%). Again, a Friedman test demonstrated a significant effect of stimulation site (*χ*^2^(2) = 24.0, *p* < 0.001). Post-hoc Wilcoxon signed-rank test with Holm correction showed significant differences between all stimulation sites (RW vs. BA, RW vs. AP, and BA vs. AP; all adjusted *p* = 0.0015), demonstrating a progressive increase in apically directed current flow from RW to BA to AP stimulation.

These findings indicate that with RW stimulation, current reaches the cochlea apex potentially exciting low-frequency fibers, however the current flowing into the apex is lower for RW stimulation than for stimulation delivered to the base (BA) or the apex (AP). In terms of acoustic masking, it is therefore possible that interaction occurs between electric stimulation delivered to the RW and low-frequency acoustic maskers. Moreover, the results show that RW stimulation produces larger longitudinal currents towards the outside of the cochlea than BA and AP stimulation, potentially increasing the likelihood of occurrence of SEs.

### 3.2. Loudness and side effects with round window, basal and apical electric stimulation

Figure 9 presents the results of the loudness level measures (T-, M-, and U-level) for each subject during stimulation via the electrode closest to the RW. Figure 9 (a) shows the measures for all tested configurations (see Figure 4 (c)-(f)), while Figure 9 (b) illustrates the loudness levels obtained for the optimized configuration selected specifically to reduce SEs with RW stimulation. If multiple configurations were able to elicit all loudness sensations, the one requiring the lowest CU was selected as optimized configuration, since higher current is associated with an increased risk of SEs (e.g., FNS; Bahmer et al., 2017). For subjects who reported SEs before reaching T-level, these events are marked with a red “x” at the corresponding CU level. An SE that was registered after the threshold is noted with the same symbol, in each case before the level that was currently targeted.

**Figure 9.**
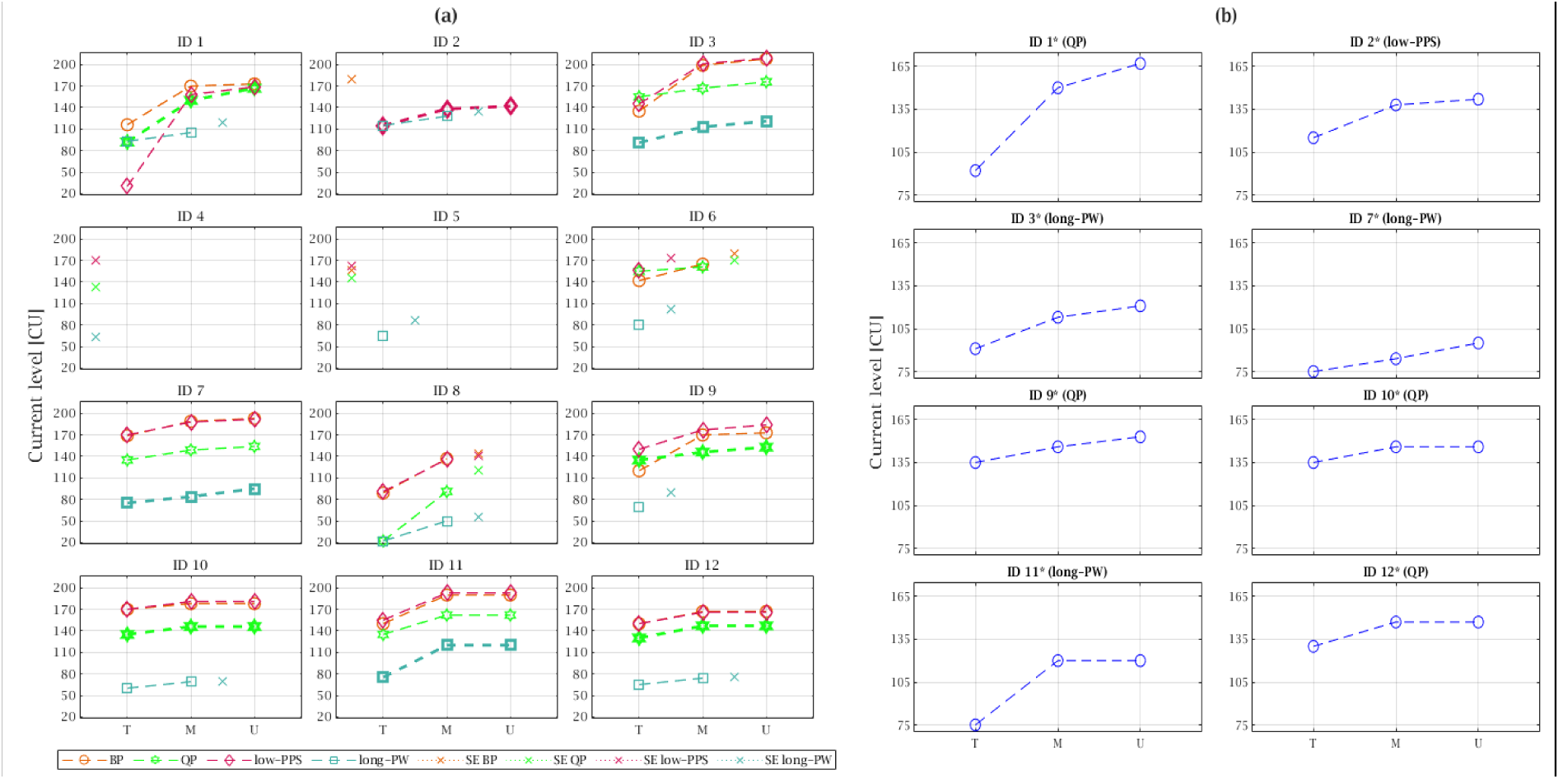
Loudness levels with electric stimulation delivered near the round window (RW), displayed in current units (CUs), for each subject. **(a)** Estimated threshold (T), most comfort (M), and uncomfortable (U) levels across all tested stimulation configurations (color-and symbol-coded). Levels at which side effects (SEs) occurred are marked by an “x” at the respective level and configuration (BP: Biphasic, QP: Quadraphasic, low-PPS: lower pulse rate, long-PW: longer phase width). **(b)** Estimated T, M, and U-levels for the individually optimized stimulation configurations to maximize loudness without SE.

Using BP (Figure 9), seven subjects (IDs 1, 3, 7, 9, 10, 11, and 12) successfully reached all three loudness levels (T, M, and U) without reporting any SEs. Two subjects (IDs 6 and 8) reached M-level but experienced SEs during stimulation (“SE BP”). Three other subjects (IDs 2, 4, and 5) reported SEs prior to any auditory perception, preventing determination of their hearing thresholds with BP stimulation.

Stimulation with quadraphasic pulses (“QP”) elicited similar loudness effects as BP stimulation. Likewise, the reduced pulse rate (“low-PPS”) configuration exhibited stimulation levels to achieve the loudness percepts, with the exception that subject ID 2 was able to perceive sound without SEs. However, two subjects (IDs 5 and 6) experienced SEs under this condition (“SE low-PPS”) and only reached the hearing threshold.

The most variable outcome among the tested configurations was observed with the longer phase width configuration (“long-PW”). Only three subjects (IDs 3, 7, and 11) reached all target loudness levels without experiencing SEs. Five subjects (IDs 1, 2, 8, 10, and 12) reached the M-level but reported SEs thereafter (“SE long-PW”), while three subjects (IDs 5, 6, and 9) only reached the hearing threshold. One subject (ID 4) neither confirmed auditory perception nor reported SEs.

An optimized configuration that met the predefined criteria was identified for eight subjects. These optimized settings varied across individuals: quadraphasic stimulation was selected for four subjects (IDs 1, 9, 10, and 12), longer phase widths were selected for three other subjects (IDs 3, 7, and 11), and one subject (ID 2) showed optimized results at a lower pulse frequency.

Figure 10 summarizes the current levels obtained when stimulating the electrode located closest to the RW for the subgroup of subjects for whom T-, M-, and U-level were measurable across the tested configurations (BP, QP, low-PPS, and long-PW). Only subjects who reported auditory perception without SEs are included, in contrast to Figure 9, which shows individual results for all participants and configurations.

**Figure 10.**
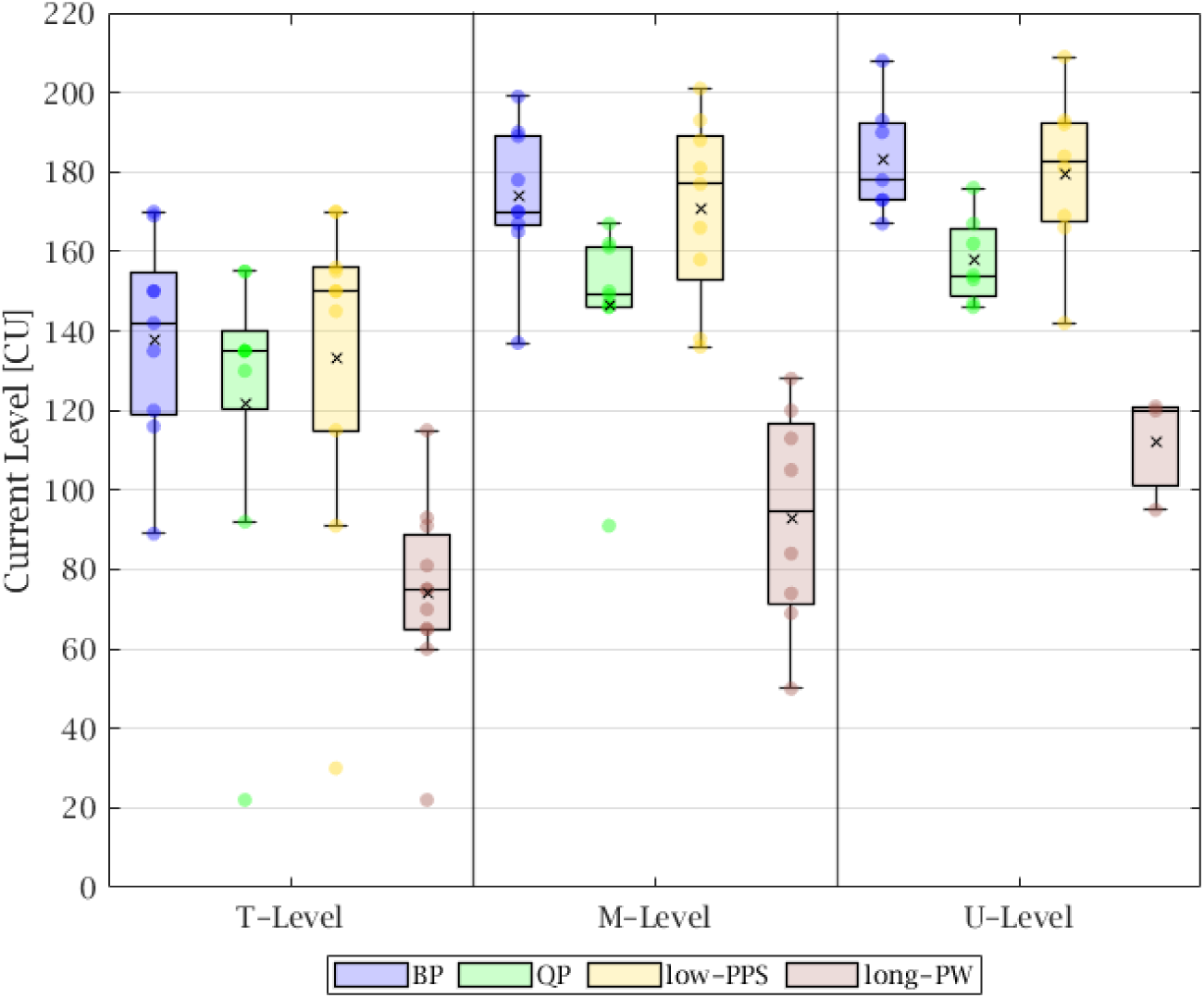
Current levels in current units (CUs) corresponding to T-, M-, and U-levels for stimulation via the electrode closest to the round window (RW), across different tested configurations: biphasic stimulation (BP, blue), quadraphasic stimulation (QP, green), reduced pulse rate (low-PPS, yellow; typically, 500 pps with individual deviations), and prolonged phase widths (long-PW, brown; 200 µs). Parameter details are listed in Table 1. Only participants who perceived auditory stimulation at the respective level without experiencing side effects (SEs) are included. Box plots display median (horizontal line), mean (cross), and the individual points in each configuration and level (circle). Individual data and additional configuration analysis are provided in Figure 9.

T-level for BP, QP, and low-PPS are comparable in CUs, whereas the long-PW configuration consistently achieves lower CU values, reflecting the effect of longer phase durations. It should be noted that CUs are not directly comparable across configurations with different phase durations in terms of total charge, as charge depends on both current amplitude and phase duration; the lower CU values observed for the long-PW configuration therefore do not necessarily indicate a lower total charge delivered. However, since current level directly determines the risk of SEs, as noted above, and since the aim of these measurements was to identify the current level at which subjects reported an auditory sensation or SEs, CUs were considered the most directly relevant parameter for this purpose. M-and U-levels diverge more strongly across configurations: BP and low-PPS present similar CUs to obtain M-level, while QP resulted in lower CUs to reach the M-level.

During stimulation, subjects experienced various SEs with differing intensities (see Table 2). The majority of SEs occurred during electrical stimulation with the RW and BA electrodes. Participants whose clinically deactivated electrodes were reactivated for the study (IDs 4, 5, and 6) reported stronger SEs more frequently. For these subjects, eliciting sound sensation via the electrode closest to the RW was not possible. Conversely, three subjects (IDs 3, 7, and 11), whose clinical maps had already the electrode closest to the RW activated, did not report any SEs.

**Table 2.** Description of side effects (SEs) reported by each participant during stimulation of the electrode closest to the round window (RW). The intensity of the SE was rated on a scale from 1 (barely noticeable) to 5 (intolerable) (see section 2.3.3. Threshold- comfort level of electric and acoustic stimuli and loudness balancing).

| Participant ID | Occurring side-effects | Strength [1 - 5] |
| --- | --- | --- |
| 1 | Dizziness/ pricking at U-Level (long-PW configuration) | Middle to strong [3 – 4] |
| 2 | Pressure at ear with stimulation at T-Level (BP configuration)<br>Selective pricking at higher electric stimulation levels (low-PPS, long-PW configuration) | Middle [3]<br>Middle [3] |
| 3 | No side-effects reported | - |
| 4 | Mostly found on forehead, no sound sensation<br>Vibration/hammering (BP-, QP-, low-PPS configuration)<br>Dizziness and nausea (long-PW configuration) | Strong [4]<br>Intolerable [5] |
| 5 | Feeling, no sound sensation (all configurations)<br>Tingling from CI/ear to the ipsilateral eye/nose (long-PW configuration)<br>For lower pps: Stitching/ pressing (low-PPS configuration) | Soft to Middle [2 – 3]<br>Middle [3] |
| 6 | No sound sensation<br>Tingling on left occiput to ipsilateral front (eye) (all configurations)<br>Drone/hammering (low-PPS configuration)<br>Feeling of electric shock (long-PW configuration) | Soft to Middle [2 – 3]<br>Middle [3]<br>Strong [4] |
| 7 | No side-effects reported | - |
| 8 | Pressure, unpleasant sensation in ear canal and forehead around U-Level (all configurations) → Also, partially at the contralateral ear side | Middle [3] |
| 9 | Unpleasant sensation in the cheek on the ipsilateral side, extending to the chin (long-PW configuration) → Pressure and pulling on the chin | Soft [2] |
| 10 | Pressure on the ipsilateral ear canal and side of the face around U-Level (long-PW configuration) | Soft [2] |
| 11 | No side-effects reported | - |
| 12 | Dizziness perceived after M-Level (long-PW configuration) | Middle [3] |

Considering the modeled current flow, the occurrence of SEs at the basal end of the cochlea can be examined in more detail (see 2.4.4. Relation between outflux current flow and side-effects). The magnitude of the current outflux (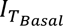, or 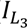 for the partially inserted participant ID 2) with RW stimulation did not appear to determine whether SEs occurred. Participants with comparable current outflux either reported SEs or did not. For example, using BP, for participants without SEs the estimated current outflux with RW stimulation were 5.13% (ID 3) and 5.78% (ID 9), while for participants with SEs the estimated current outflux was similar of 7.07% for ID 4 and 9.87% for ID 5.

### 3.3. Dynamic Range

Dynamic ranges (DRs) were defined as the difference between M-and T-levels in CUs. T-levels across stimulation configurations (BP, QP, low-PPS, long-PW) and electrode positions (RW, BA, AP) were obtained through the 3-AFC procedure.

In BP (Figure 11 (a)), DRs were lower for BA stimulation (median 25.5 CU, range 11.5–52.8 CU) compared to AP stimulation (median 43.75 CU, range 14–97.5 CU). In the optimized configuration (Figure 11 (b)), analyzed in the subgroup of eight subjects who perceived all three loudness levels with RW stimulation without SEs (see 2.3.3. Threshold-comfort level of electric and acoustic stimuli and loudness balancing), RW stimulation yielded the lowest DR (median 19.5 CU, range 11.5–68.3 CU), followed by BA (median 33.05 CU, range 14.3–50.8 CU) and AP (median 55.4 CU, range 17.5–75 CU). The difference between BA and AP was not statistically significant.

**Figure 11.**
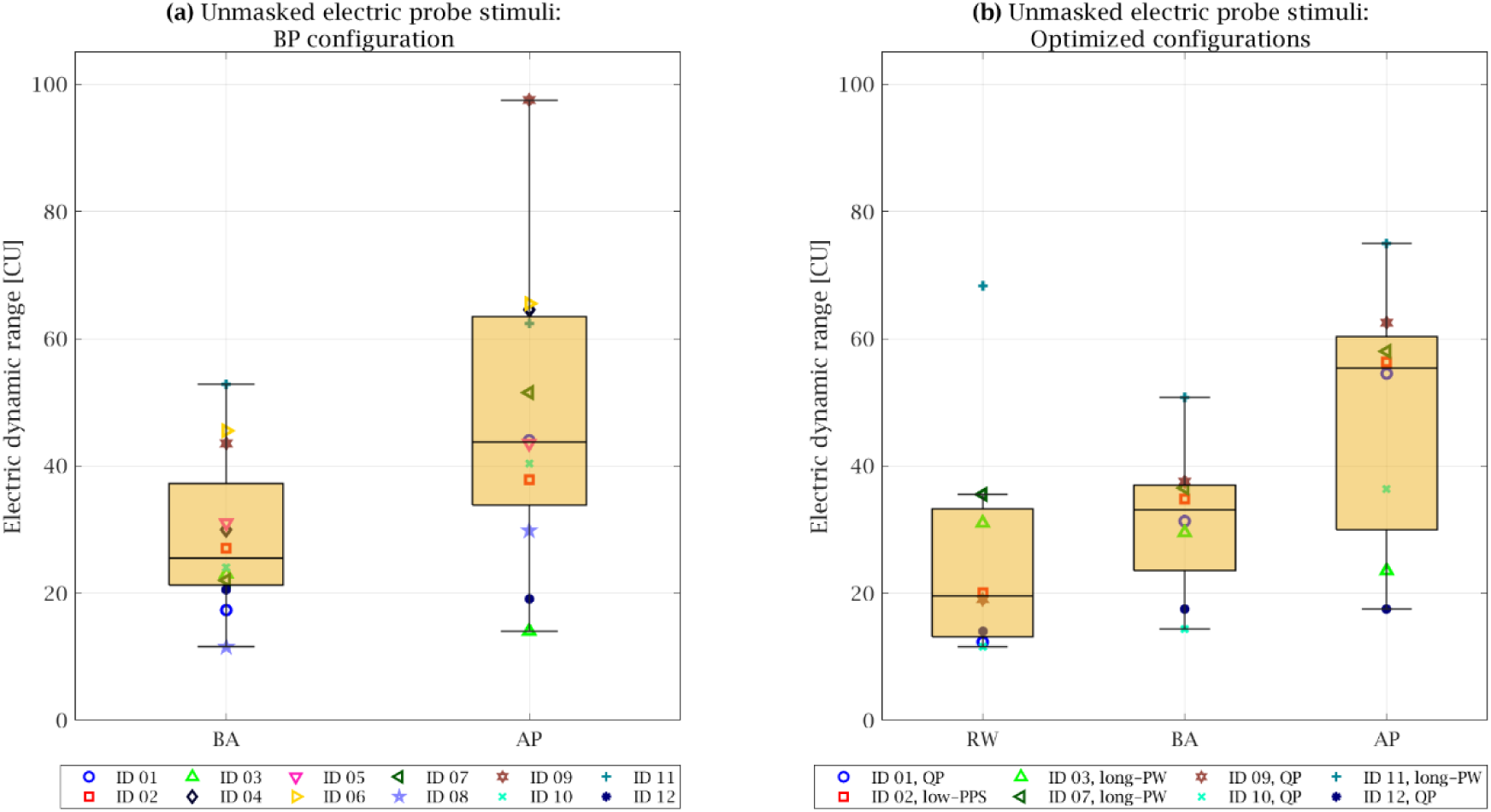
Electrical dynamic range across subjects for unmasked electrical probes at different electrodes. **(a)** The dynamic ranges measured under the biphasic stimulation (BP), which were observed in all subjects. Stimulation was applied using the respective basal (BA) and apical (AP) electrodes. **(b)** Dynamic ranges for the optimized configuration of electric stimulation, determined for eight subjects (see Figure 9 (b)). In this case, stimulation included the round window (RW) electrode in addition. The optimized stimulation configuration for each subject is indicated in the legend, while individual subjects are distinguished by color and marker symbol.

### 3.4. Masking Results

Figure 12 shows the TE caused by acoustic maskers on electric stimulation with BP across all 12 subjects, with stimulation at the BA and AP electrodes. TEs are shown in CU for *fa1* (250 Hz; top), *fa2* (individual higher frequencies; middle), and *GN* (broadband WB-noise; bottom). The corresponding right panels display the same data as a percentage of the electrical dynamic range (%DR). TEs in CU ranged from –6 CU (ID 1; WB-noise; AP) to 29 CU (ID 9; WB-noise; AP). TEs expressed as percentage of DR (%DR), values ranged from –13.6% (ID 1; WB-noise; AP) to 56.5% (ID 8; 1 kHz; BA).

**Figure 12.**
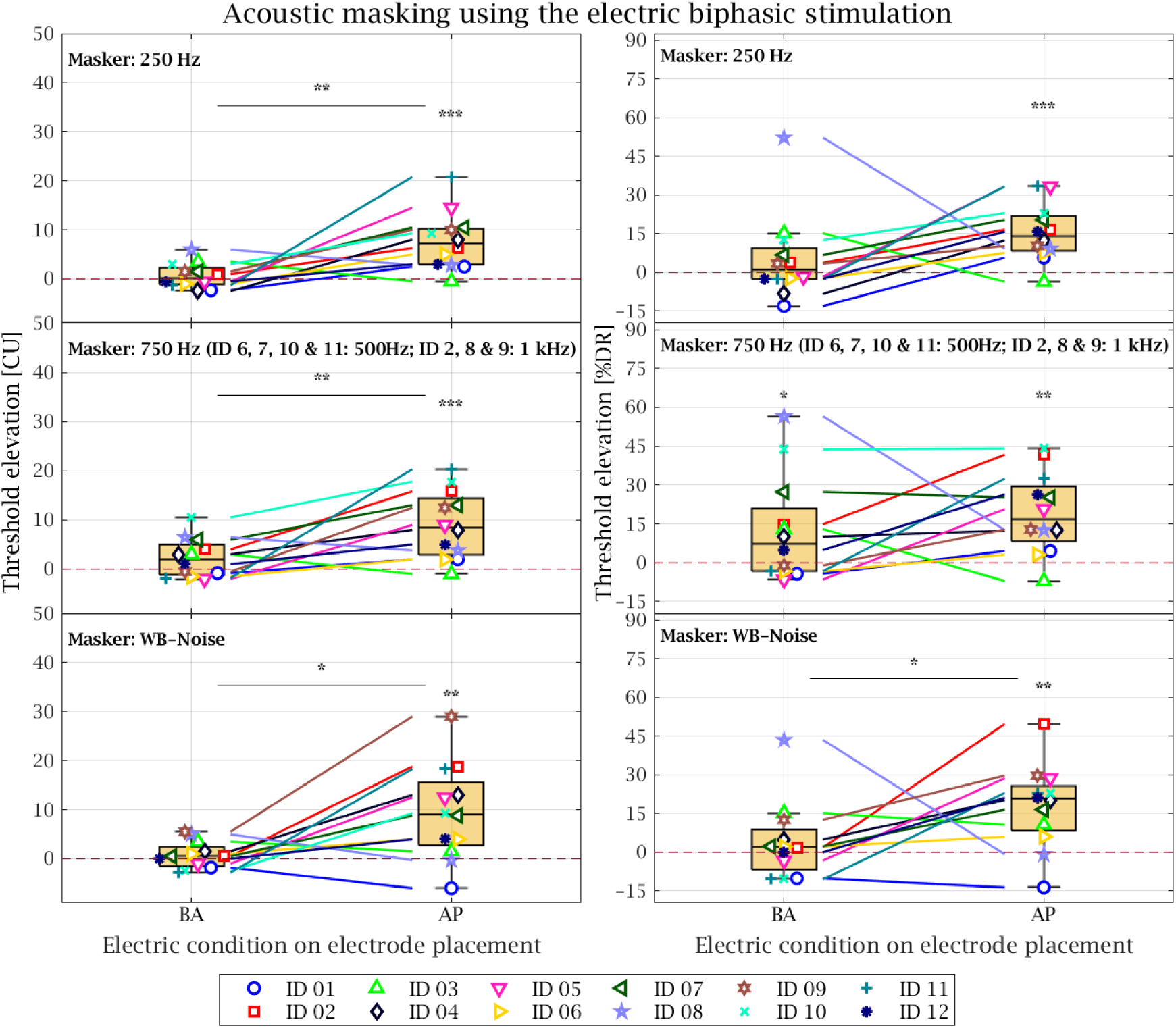
Acoustic masking of electrical probes using the clinical electrical stimulation setting (biphasic stimulation; BP) for each subject (see Table 1) expressed as threshold elevation (TE). Left panels display TEs in current units (CU), while right panels show TEs as a percentage of the dynamic range (%DR). Individual subjects are identified by unique combinations of color and marker symbol. The stimulation electrodes are shown on the x-axis as the basal active (BA) and most apical (AP) electrodes. Each panel corresponds to a different acoustic masking condition: Top – pure tones at *fa1* (250 Hz); Middle – pure tones at *fa2* (750 Hz, except for subjects ID 6, 7, 10, 11 at 500 Hz, and ID 2, 8, 9 at 1 kHz); Bottom – *GN* (broadband white-band noise; WB). Lines connecting BA and AP electrodes indicate an increase or decrease in TE. Asterisks mark statistically significant TE (*p* < 0.05, Wilcoxon rank-sum test; Table 3).

**Table 3.** Threshold elevations (TE) for each electrode, given in current units (CU) and dynamic ranges (%DR), with corresponding ***p*** values (Wilcoxon signed-rank test) for acoustic masking under biphasic and optimized stimulation configurations. Abbreviations: RW: round window electrode; BA: basal active electrode; AP: most apical electrode; BA–RW: difference between BA and RW; AP–RW: difference between AP and RW; AP–BS: difference between AP and BA.

| Masker | Electrode | Acoustic Masking with Biphasic Configuration |  |  |  | Acoustic Masking with Optimized Configurations |  |  |  |
| --- | --- | --- | --- | --- | --- | --- | --- | --- | --- |
|  |  | Mean TE [CU] | *p* | Mean TE [%DR] | *p* | Mean TE [CU] | *p* | Mean TE [%DR] | *p* |
| *f*<sub>a1</sub><br>(250 Hz) | RW | - | - | - | - | 1.0 | 0.453 | 2.8 | 0.344 |
|  | BA | 0.7 | 0.218 | 5.3 | 0.206 | 1.5 | 0.227 | 6.5 | 0.234 |
|  | AP | **7.7** | **<0.001** | **15.3** | **<0.001** | **7.1** | **0.016** | **16.0** | **0.012** |
|  | BA–RW | - | - | - | - | 0.5 | 0.219 | 3.7 | 0.289 |
|  | AP–RW | - | - | - | - | 6.1 | 0.059 | 13.2 | 0.098 |
|  | AP–BA | **7.0** | **0.002** | 10.0 | 0.076 | 5.6 | 0.156 | 9.5 | 0.191 |
| *f*<sub>a2</sub><br>(500 Hz,<br>750 Hz,<br>1000 Hz) | RW | - | - | - | - | 3.0 | 0.172 | 14.6 | 0.109 |
|  | BA | 2.3 | 0.053 | 12.7 | **0.037** | 0.2 | 0.438 | 2.3 | 0.422 |
|  | AP | **9.0** | **<0.001** | **19.1** | **0.001** | **10.9** | **0.016** | **22.1** | **0.016** |
|  | BA–RW | - | - | - | - | -2.8 | 0.984 | -12.3 | 0.984 |
|  | AP–RW | - | - | - | - | 7.9 | 0.219 | 7.5 | 0.422 |
|  | AP–BA | **6.7** | **0.003** | 6.4 | 0.088 | 10.7 | 0.078 | 19.8 | 0.078 |
| *GN*<br>(White-Band<br>Noise) | RW | - | - | - | - | 2.2 | 0.172 | 1.6 | 0.406 |
|  | BA | 0.8 | 0.239 | 4.1 | 0.252 | 2.5 | 0.5 | 9.4 | 0.094 |
|  | AP | **9.4** | **0.003** | **17.9** | **0.002** | **11.3** | **0.016** | **24.8** | **0.016** |
|  | BA–RW | - | - | - | - | 0.3 | 0.578 | 7.8 | 0.203 |
|  | AP–RW | - | - | - | - | 9.1 | 0.109 | 23.2 | 0.094 |
|  | AP–BA | **8.6** | **0.011** | **13.8** | **0.031** | 8.8 | 0.156 | 15.4 | 0.156 |

A Wilcoxon signed-rank test (Table 3) revealed significant TEs for AP stimulation across all acoustic maskers (*fa1*, *fa2* and *GN*). For BA stimulation, a significant elevation was only observed for higher-frequency maskers (*fa2*) when expressed in %DR (*p* = 0.037). Comparisons between BA and AP electrodes showed significantly higher TEs at the AP for all maskers in CU (250 Hz: *p* = 0.002; higher frequency: *p* = 0.003; WB-noise: *p* = 0.011), and for WB-noise in %DR (*p* = 0.031). Considerable inter-individual variability was observed. Several subjects (notably ID 3 and 8; to a lesser extent ID 1, 7, and 10) showed lower TEs with AP stimulation compared to BA, particularly in %DR (Figure 12, right).

Figure 13 illustrates the TEs observed in the acoustic masking experiment under optimized stimulation configurations. The electric thresholds ranged from –12.3 CU (ID 11; masker: 500 Hz; long-PW with BA) to 35.8 CU (ID 2; masker: WB-noise; low-PPS with AP). Expressed as %DR, TEs varied between –26.1% DR (ID 10; probe: QP with RW; masker: WB-noise) and 63.6% DR (ID 2; probe: low-PPS with AP; masker: WB-noise).

**Figure 13.**
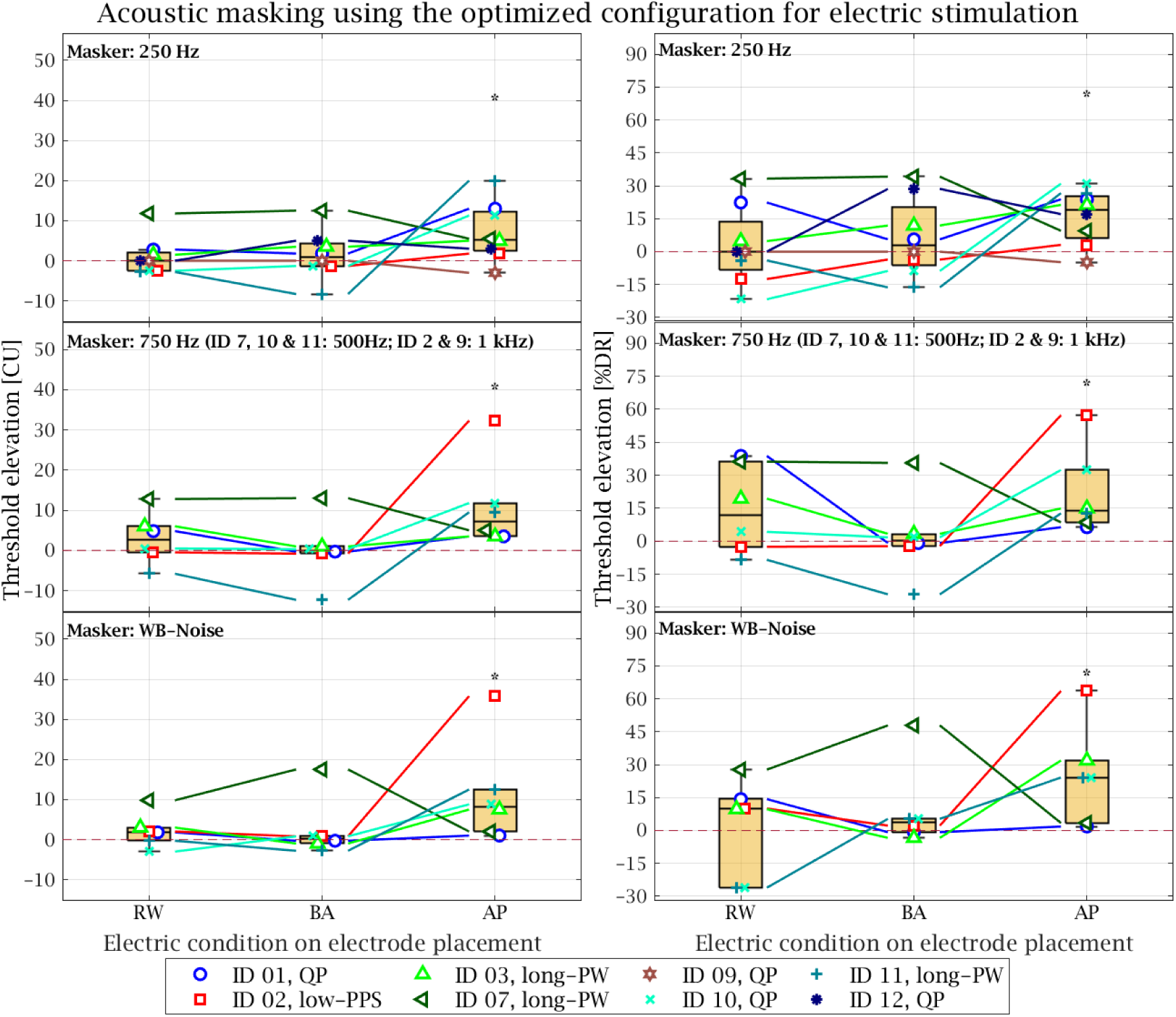
Acoustic masking of electrical probes using the subject-specific optimized electrical stimulation settings (see Table 1), expressed as threshold elevation (TE). Left panels show TEs in current units (CU), right panels as a percentage of the dynamic range (%DR). The optimized stimulation configuration for each subject is indicated in the legend, while individual subjects are distinguished by color and marker symbol. The stimulating electrodes is displayed on the x-axis: round window (RW), basal (BA), and apical (AP). Panels correspond to masker types: Top – pure tones at *fa1* (250 Hz); Middle – pure tones at *fa2* (750 Hz; except ID 7, 10, 11: 500 Hz; ID 2, 9: 1 kHz); Bottom – *GN* (broadband white-band noise; WB). Connecting lines indicate changes in TE across stimulating electrodes. Asterisks mark statistically significant effects (***p*** < 0.05, Wilcoxon signed-rank test, see Table 3).

Significant TEs were observed for AP electrode stimulation for both when expressed in CU and %DR, as shown in Table 3 (right panel). Overall, observed TEs revealed high individual variability. Some subjects (IDs 1, 3, 7, and 12) exhibited masking when electric stimulation was provided to the RW and BA electrode with their optimized configurations. Notably, Subject ID 7 showed pronounced TEs with RW and BA electrode stimulation across all acoustic maskers, but not with AP stimulation. In contrast, Subject ID 2 displayed significant TEs with AP stimulation compared to BA and RW stimulation, particularly with the higher-frequency (*fa2*) and WB-noise (*GN*) maskers.

### 3.5 Effect of hearing loss on acoustic masking

The relation between TE and residual hearing was investigated through correlation analyses. Averaged TEs expressed in CU for acoustic maskers between *fa1* (250 Hz) and *fa2* (e.g., 750 Hz) showed a significant positive correlation with PTA thresholds averaged across 125–2000 Hz (Figure 14 (c); *p* = 0.024). A positive but non-significant trend was observed for higher-frequency (*fa2*) maskers (Figure 14 (b); *p* = 0.053). In contrast, no correlation was found between TE induced by a 250 Hz (*fa1*) masker and the corresponding 250 Hz PTA (Figure 14 (a); *p* = 0.509).

**Figure 14.**
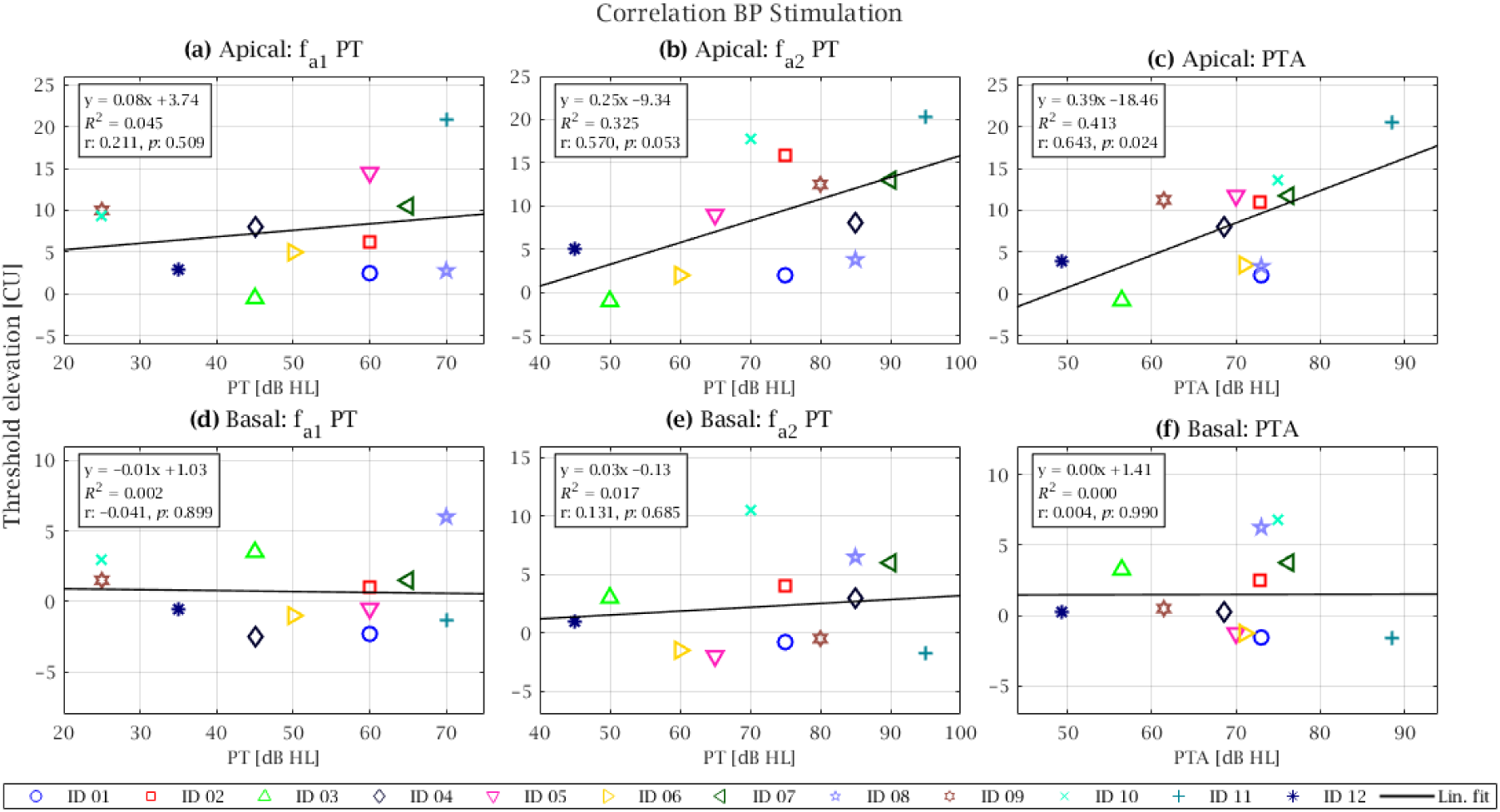
Correlation analysis between threshold elevations (TEs) and audiometric pure-tone (PT) measures for the biphasic (BP) stimulation configuration. **Panels (a–c)** show correlations for apical (AP) electrical stimulation, and **panels (d***–***f)** for basal (BA) stimulation. Individual subjects are identified by unique combinations of color and marker symbol. For each stimulation site, correlations are shown between TEs induced by the low-frequency masker *fa1* (250 Hz) and the corresponding audiometric threshold **(a, d)**, between TEs induced by the individual higher-frequency masker *fa2* and the corresponding audiometric threshold **(b, e)**, and between the mean TE across *fa1* and *fa2* maskers and the pure-tone average (PTA; 125-2000 Hz) **(c, f)**. Solid lines indicate linear fits; Pearson’s *R*² and slope values are reported in each panel.

When TEs were normalized to the individual electrical dynamic range (%DR), no significant correlations with residual hearing were observed for any masker frequency or stimulation site.

Correlation analyses were also performed for optimized stimulation settings as well as for basal and RW stimulation (see Figure 15). However, none of these conditions showed significant correlations between TE and residual hearing; therefore, subsequent analyses focused on biphasic (BP) stimulation at the apical electrode.

**Figure 15.**
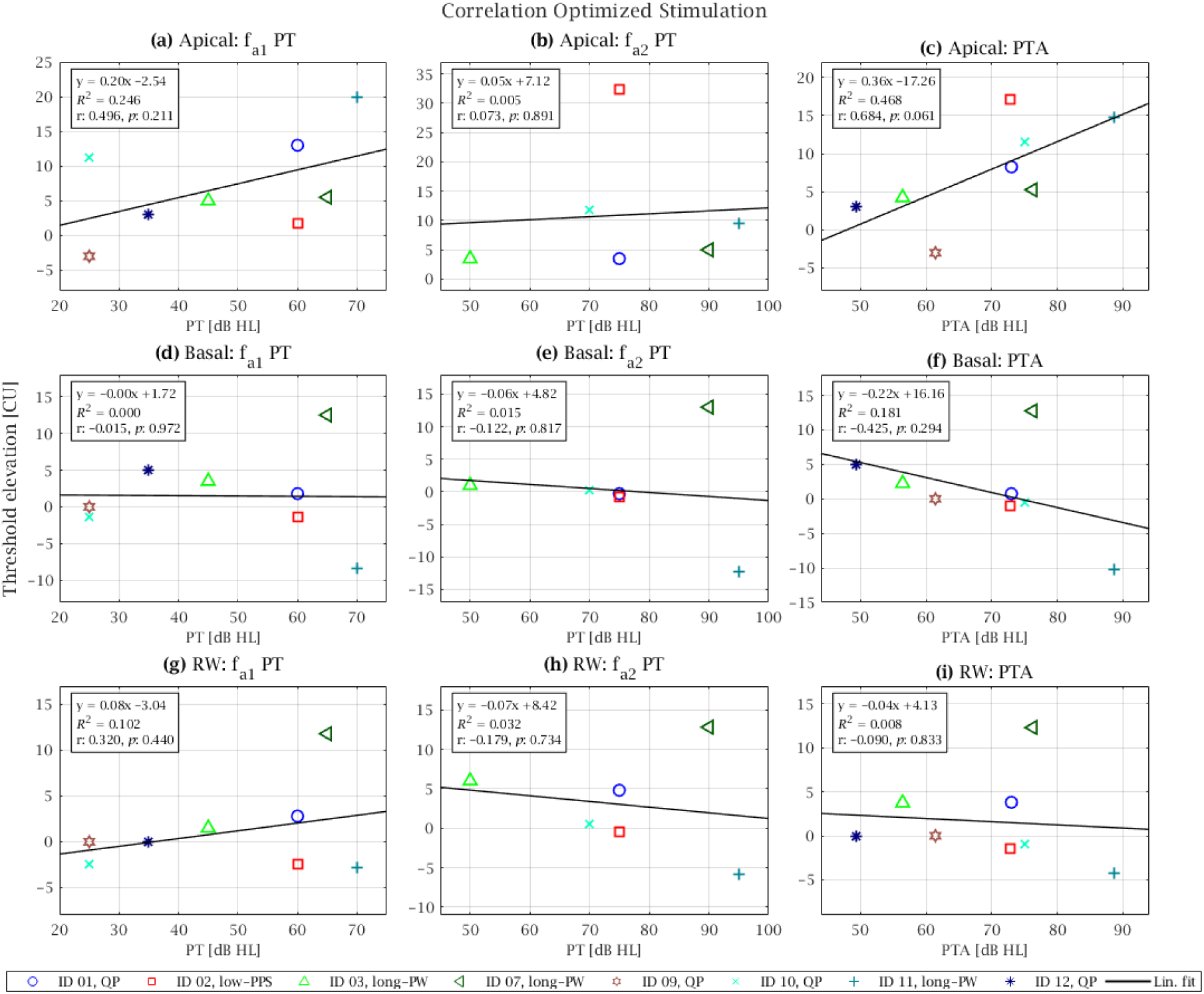
Correlation analysis between threshold elevations (TEs) and audiometric pure-tone (PT) measures for the optimized stimulation configurations. **Panels (a***–***c)** show correlations for apical (AP) electrical stimulation, **panels (d***–***f)** for basal (BA) stimulation, and **panels (g***–***i)** for round window (RW) stimulation. The optimized stimulation configuration for each subject is indicated in the legend, while individual subjects are distinguished by color and marker symbol. For each stimulation site, correlations are shown between TEs induced by the low-frequency masker *fa1* (250 Hz) and the corresponding audiometric threshold **(a, d, g)**, between TEs induced by the individual higher-frequency masker *fa2* and the corresponding audiometric threshold **(b, e, h)**, and between the mean TE across *fa1* and *fa2* maskers and the pure-tone average (PTA; 125-2000 Hz) **(c, f, i)**. Solid lines indicate linear fits; Pearson’s *R*² and slope values are reported in each panel.

## 4. Discussion

This study investigated simultaneous ipsilateral acoustic masking of electrical stimulation delivered near the RW by low-frequency acoustic tones in twelve EAS users. Acoustic masking was compared across electric stimulation sites, including electrical stimulation near the RW, within the basal region, and at the most apical intra-cochlear electrode. In addition, the influence of different electric stimulation configurations applied near the RW on loudness and SEs was assessed.

This study examined whether electrical stimulation at the cochlear base or RW can be acoustically masked and whether this effect can be used to assess residual low-frequency hearing, while also considering the potential undesirable SEs associated with basal and RW stimulation.

Across configurations, acoustic masking of electrical stimulation was consistently observed. Masking was most pronounced for apical stimulation and markedly reduced when electric stimulation was delivered at the base or near the RW, thereby confirming and extending previous findings. The results further showed that acoustic masking was associated with hearing loss when electrical stimulation was delivered to the most apical electrode.

To explore potential mechanisms underlying these observations, a lumped-parameter model of intra-cochlear current flow was employed. The model suggests that electrical stimulation may spread toward the apex and activate low-frequency auditory nerve fibers, which are subsequently masked by low-frequency acoustic stimulation at the nerve.

Taken together, these findings provide new insight into the interaction between acoustic and electric stimulation in EAS users. Acoustic masking of apical electric stimulation, which correlated with residual hearing, suggests potential for clinical assessment of residual low-frequency hearing on a post-operative basis. In contrast, masking of basal and RW stimulation was consistently measurable but did not correlate with residual hearing, and its clinical application as a pre-operative diagnostic measure therefore needs further studies to assess its possible application, particularly given the substantial inter-subject variability observed, the relatively high incidence of stimulation-related side effects, and the practical challenges of conducting psychophysical measurements of this kind in young children. Nonetheless, these findings provide novel insight into auditory perception and side effects associated with basal and extra-cochlear electric stimulation, which may inform future efforts to optimize this approach, for example through improved electrode placement or stimulation parameters, before its clinical utility as a diagnostic tool can be more conclusively established.

### 4.1. Trans-impedance measurements and simulated cochlear current flow

Trans-impedances recorded during RW stimulation demonstrated that electrical current flows inside the cochlea (Figure 6), confirming that RW stimulation can engage cochlear structures and thereby interact with acoustic input. Simulated current-flow patterns from the lumped-parameter model (Figure 7 (a), (c), (e)), showed that RW and basal intra-cochlear stimulation produced a larger proportion of current leaving the cochlea than apical stimulation (Figure 8 (a), median extra-cochlear current: 17.2% for RW vs. 8.4% for apical stimulation). This indicates that basal and RW stimulation drive current toward extra-cochlear pathways, increasing the likelihood of current spread to adjacent non-auditory structures, such as the tympanic segment of the FN (Saoji et al., 2024). Consistent with this, longitudinal current components (Figure 7 (b), (d), (f)) showed that both RW and basal stimulation leaked current into the extra-cochlear space. In subject ID 2, current levels dropped particularly steeply once leaving the cochlea (Figure 7 (b), (d)), suggesting that this extra-cochlear flow could activate surrounding tissue or neural structures and thereby contribute to SEs.

Notably, both RW and BA stimulation also produced transversal current directed toward the apical ST (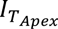 Figure 7 (a), (c); Figure 8 (b)), followed by a non-monotonic, reciprocal pattern: high near the stimulation site, decreasing toward mid-cochlearly, and rising again toward the most apical contacts (see also the schematic in Figure 5). This pattern reflects the topology of the lumped-parameter model (Vanpoucke et al., 2004) rather than a distinct anatomical exit route at the apex: the modeled ST is terminated basally by a resistor representing extra-cochlear pathways (e.g., cochlear and vestibular aqueducts, internal auditory canal, middle ear), capturing much of the current near any basal stimulation site, whereas the apical end has no equivalent shunt – the most apical node is a dead end, forcing any current that reaches it to exit transversally there. This explains why apical electrode stimulation itself (Figure 7 (e)) does not show this dip-then-rise pattern, since the injection site is already close to the apical terminal node. Because the modeled *IT* is derived from the fitted trans-impedance profile as a whole rather than the local impedance at a single contact, it need not correspond to a measurable impedance change at the apical electrodes themselves. For comparison, apical electrode stimulation produced the highest overall proportion of apically directed transversal current (see Figure 8 (b)), consistent with its proximity to the apical turn.

Together, these findings indicate that extra-cochlear RW (and basal) stimulation can reach low-frequency cochlear regions, providing a plausible mechanism by which such stimulation activates low-frequency neural populations. These populations are also engaged by low-frequency acoustic stimuli, which may drive them into a refractory state and thereby mask the electrical stimulation delivered through the RW – contributing to the TEs observed during acoustic masking.

Finally, the current-flow model was designed to capture the dominant pathways of intra-and extra-cochlear current spread relevant to present research question. Real current trajectories in vivo may include additional pathways not captured here, but these would primarily affect fine-grained spatial details rather than overall distribution trends identified. The key findings – apically directed current during RW/basal stimulation and increased extra-cochlear current during basal stimulation – remain robust within this modeling framework. Future refinements may improve spatial resolution but are not required to support the present mechanistic interpretations.

### 4.2. Loudness with electric stimulation delivered around the round window

Loudness was measured by delivering current through a CI electrode positioned as close as possible to the RW. The present study demonstrates that auditory perception through electrical stimulation of this electrode is feasible without unwanted SEs. However, the strength and nature of SEs varied considerably across subjects (see Table 2).

Not all stimulation configurations elicited an auditory percept in every subject, underlining interindividual variability. Threshold differences across configurations highlight the importance of tailored stimulation parameters, such as phase width and pulse rate, to optimize auditory perception while minimizing SEs.

Subjects for whom the RW electrode was deactivated in their clinical program (ID 4, 5, 6, 11, 12) were more likely to report SEs when RW electrode was activated. These effects intensified with increasing current at the RW electrode, limiting the perceived loudness. The timing of clinical deactivation varied across these subjects: for three subjects (ID 4, 5, 12), the RW electrode was already deactivated at the first clinical fitting, indicating that SEs or poor auditory perception were present from the onset of CI use. This finding suggests that these issues were more likely related to the anatomical position of the electrode relative to sensitive structures, such as the FN, rather than a progressive tissue response. For one subject (ID 11), deactivation occurred two months after first fitting, and for another (ID 6), only after approximately one year; for the latter case, the delayed onset is more consistent with the time course of fibrotic encapsulation around the electrode, which has been suggested as a possible contributor to increased impedance and altered current spread despite trans-impedances suggesting technically functional operation (Saoji et al., 2024). In some subjects (e.g., ID 4 and 5), SEs could not be altered or overcome, even when different configurations of electric stimulation were tested. Several subjects also experienced difficulty distinguishing between auditory perception and general somatosensory sensations, as the stimulation felt unfamiliar. This further reduced the likelihood of identifying an optimized set of basal stimulation parameters.

In contrast, in other participants (ID 3, 7, 8, 11), the choice of electric stimulation configuration (BP, QP, low-PPS, long-PW) had little influence on perception, as they consistently reported comparable auditory thresholds when the stimulation was delivered close to the RW. As shown in Figure 9, subjects IDs 3, 7, and 11 achieved T-, M-, and U-levels without SEs, while subject ID 8 reached T-and M-levels but not U-level. For these subjects, results were stable across all tested configurations. The remaining subjects (ID 1, 2, 9, 10, 12) exhibited configuration-dependent outcomes, with some settings producing auditory threshold sensations and other settings eliciting SEs. This suggests that individual selection of stimulation parameters may be critical for obtaining T-, M-, and U-levels when electric stimulation is delivered at the RW electrode.

Previous work demonstrated that extending phase widths can reduce FNS by lowering current amplitude (Alahmadi et al., 2023; Gärtner et al., 2023). Consistent with this result, the long-PW applied to the RW electrode resulted in diminished SEs as lower T-, M-, and U-levels were required, see for example subjects with IDs 3, 7, and 11 in Figure 10.

Still, some subjects experienced SEs with the longer-PW condition (Figure 9). Earlier studies reported that TP stimulation typically requires stimulation higher amplitude (charge) than BP stimulation to achieve equal loudness when total phase duration is constant, yet it may reduce unwanted FNS, likely due to altered neural activation patterns compared with BP pulses (Bahmer et al., 2017; Braun et al., 2019). In the current study, we used quadraphasic (pseudo-triphasic) pulses having a total duration twice as the BP pulses to assess if quadraphasic can reduce SEs. On average, this configuration required ∼20 CU less current than BP stimulation. This reduction was not formally tested for statistical significance but is descriptively in the same direction as predicted by charge-based loudness constancy: doubling phase duration while maintaining equal charge would theoretically allow current to be halved (a 6 dB reduction), which corresponds to approximately 38 CU given the current-to-CU conversion used by the stimulation system (see 2.2.2 Electric stimuli). The smaller reduction observed (∼20 CU) may thus be explained by the QP pulse shape being less efficient than the BP pulse shape. In four subjects (ID 1, 9, 10, 12), QP stimulation reduced SEs, though this benefit may have been partly due to the longer phase duration. Besides, QP requires more current than longer-PW. On average, the longer-PW requires ∼50 CU less current than QP. In two participants (ID 2, 6), QP stimulation either increased SEs or provided no advantage compared to other configurations (BP, low-PPS, long-PW; Figure 9). Reducing the pulse rate (low-PPS condition) did not have an impact on reducing SEs. On average, the current required for T-, M-, or U-levels at low-PPS was similar to that with BP stimulation, only one subject reported reduced SEs with this configuration.

Beyond stimulation settings, electrode placement itself can influence the occurrence of SEs. For example, electrodes positioned close to the facial nerve may inadvertently activate it even at low current levels (Saoji et al., 2024). Suboptimal electrode positioning may also affect other outcome measures, including masking results, which are discussed in more detail in a later section. Uncertainties related to CBCT-based assessment of electrode position may further contribute to inter-subject variability (Thormählen et al., 2024). Pre-existing conditions such as tinnitus (as observed in subject ID 1) can further complicate the process of determining suitable stimulation parameters (Kheirkhah et al., 2024). In summary, adapting stimulation configurations (e.g., QP, low-PPS, long-PW) when delivering current close to the RW may help reduce SEs; however, effects appear to be highly subject-specific.

### 4.3. Electric dynamic range of electric stimuli delivered at different locations of the cochlea

Electric DR provides important context for interpreting acoustic masking results expressed in CU and as %DR. Across subjects, electrical stimulation delivered to the basal region of the cochlea, including stimulation near the RW, consistently produced a smaller DR compared to stimulation at more apical electrode sites, for both BP and optimized configurations (Figure 11). Importantly, this demonstrates that RW and basal stimulation reliably support graded loudness perception across multiple stimulation levels, confirming their suitability for perceptual and masking experiments. The observed differences in DR across stimulation sites are consistent with previous reports describing larger DRs when electric stimulation is provided at more apical electrodes (Kipping et al., 2020; Krüger et al., 2017).

In the context of acoustic masking, these location-dependent DR characteristics have implications. At basal and RW stimulation sites, the available loudness range is more reduced, which naturally narrows the range over which TEs can be expressed. As a result, masking can still be observed at these locations, but their quantification is inherently less granular than at apical sites, where a larger DR allows finer resolution of threshold changes. Accordingly, the more pronounced and consistently quantifiable masking observed at apical electrodes can be attributed, at least in part, to these intrinsic DR characteristics rather than to fundamental differences in the underlying masking mechanisms.

### 4.4. Acoustic masking of electric stimulation

Regardless of whether the acoustic masker consisted of pure tones or white noise, it caused a TE of electric stimulation delivered to the apical electrode, confirming the findings of Krüger et al. (2017), Imsiecke et al. (2020), and Kipping et al. (2020). Acoustic masking of electric probes was significantly lower when stimulation was delivered to the basal active electrode compared to the apical electrode (*p* < 0.05).

In Krüger et al. (2017), acoustic masking was observed from the most apical electrode (EL22) to the most basal electrode used in that study (EL17; in ID3: EL20). Masking was present when acoustic stimulation consisted of a 250 Hz pure tone (–0.5 CU; –1.5% DR), or a pure tone with a frequency around 750 Hz (–0.5 CU; –10% DR), and when TEs were averaged across all acoustically tested frequencies available for each subject. Depending on the subject, these frequencies covered a range from 125 Hz up to 1500 Hz (as shown in Fig. 8 in Krüger et al., 2017). The corresponding mean masking across this full measured range was –0.3 CU; –15% DR).

#### 4.4.1. Biphasic stimulation

Acoustic masking caused significant TE of electrical probes in the BP configuration. The strongest masking effects occurred for stimulation at the most apical electrode, with a mean TE of 17.4 %DR (8.7 CU). These findings are in close agreement with previous works by Krüger et al. (2017), who reported a mean TE of 21.5 %DR (9.5 CU) with a maximum of 55.2 %DR (26.5 CU) in apical electrodes and Kipping et al. (2020), who found a mean TE of 18.7 %DR (8.6 CU) and a maximum of 59 %DR (23.8 CU) in Hybrid-L users.

Across subjects, and with few exceptions (ID 1, 3, and 8), TEs were consistently higher for apical than for basal stimulation sites (Figure 12). Therefore, these results support that apical stimulation is more susceptible to acoustic masking. This is consistent with previous work showing that TE patterns between apical (EL22/EL21) and more medial electrodes (EL17) vary across individuals but typically exhibit larger TE at apical sites (Kipping et al., 2020; Krüger et al., 2017). In line with Kipping et al. (2020), the higher-frequency masker (e.g., 750 Hz) in the present study produced a significant basal-apical TE difference when expressed as %DR.

Importantly, acoustic masking was not limited to apical stimulation. All masker types, including both tonal (*fa1* and *fa2*) and broadband (*GN*) stimuli, produced measurable TEs at the basal stimulation site. While TEs expressed in CU were significantly higher at the apical electrode, basal stimulation still exhibited detectable masking, demonstrating that acoustic masking is possible even when electrical stimulation is delivered close to the RW or the base of the cochlea. When TEs were expressed as %DR, the basal-to-apical difference remained significant primarily for the WB-noise masker, indicating normalization of DR affects the strength of frequency-specific masking.

For basal stimulation, a significant TE was observed mainly for a higher-frequency (*fa2*) masker, and only in %DR. Rather than indicating an absence of masking, this finding highlights that acoustic masking at basal sites is smaller in magnitude and therefore detectable only under specific acoustic and stimulation conditions. Crucially, these results provide the first systematic evidence that acoustic masking of electrically evoked percepts can occur even when stimulation is delivered near the RW or the cochlear base, which is central to the aims of the present study.

The stronger masking observed for the higher-frequency masker is consistent with the closer spatial correspondence between the acoustic stimulation frequency (e.g., 750 Hz) and the region activated by basal electrical stimulation. This spatial proximity, together with the modeled transversal current components toward apical cochlear regions, likely facilitates increased interaction between acoustic and electric excitation. As such, higher-frequency acoustic stimulation may be particularly suitable for revealing residual hearing when electric stimulation is delivered at basal sites, provided that sufficient acoustic sensitivity remains in the corresponding frequency range.

Finally, marked inter-individual variability was evident. For example, in subject ID 9, WB-noise produced larger TE than tonal maskers (*fa1* and *fa2*) at both basal and apical electrodes, underscoring that acoustic masking in EAS is highly subject-specific and influenced by individual auditory and cochlear factors.

#### 4.4.2. Optimized configurations

Different stimulation configurations were investigated to increase auditory sensations while minimizing SEs. Similar to the TE in the BP configuration (see 4.4.1. Biphasic stimulation), significant TE was observed for the most apical electrode when stimulating with the respective optimized configuration, but not for the basal electrodes (RW and BA). The mean TE for the apical electrode under the optimized configurations is 21.0 %DR (9.8 CU).

Interestingly, a stronger individual TE, particularly in %DR, was observed when stimulating the RW electrode under the presence of a higher-frequency masker, even though this effect did not reach significance at the group level. This resembles the findings for the BP configuration, where the higher-frequency masker produced a significant TE difference (see 4.4.1. Biphasic stimulation). In addition, for some individuals an increase in TE was also evident with the lower-frequency masker, which had not been observed under the clinically based BP configuration. This suggests that, with optimized stimulation in the basal region and a low-frequency masker, TE might be used to assess residual hearing more effectively. Furthermore, some subjects (e.g., ID 2 with high-frequency and WB noise maskers) showed comparatively high TEs.

Due to the large inter-individual variability in TE, steeper linear increases and decreases between electrodes are apparent, particularly with the higher-frequency masker (see Figure 13). In Kipping et al. (2020), several configurations were tested in addition to the clinical BP configuration. With one of these configurations, TE was measured at apical (EL22 and EL21) and medial (EL17) electrodes. Depending on the acoustic masker, TE from apical to medial stimulation either increased or decreased, indicating that TE in the basal region is highly subject-specific. Similar patterns were observed in the present study: some subjects (e.g., ID 2 and 7 at most apical electrode) displayed a pronounced increase or decrease in TE across electrodes. However, because these effects were confined to individual cases and occurred across different configurations, no generalizable conclusion can be drawn.

Regarding stimulation in the basal region (RW and BA electrode), masking appears to be individual, similar to those observed under BP. Acoustic masking in this region was minimally affected by the low-PPS (shorter pulse rate) and the QP (quadraphasic) configurations, except in individual cases such as subjects ID 1 and 12. In contrast, with the long-PW configuration, TE showed a tendency to increase in subject ID 7 and, to a lesser extent, in subject ID 3. Additionally, the choice of masker generally did not influence TE, except in subjects ID 2 and 7 for the apical electrode, where clear masker-dependent differences were observed. Individual differences in TE may be explained by the presence of multiple neural pathways and interaction mechanisms. One manifestation of this individual variability is that TEs were not uniformly positive across subjects, electrodes, and masker conditions: at the group level, TE was tested for significance against zero, but at the individual level only descriptive observations can be made. Several subjects showed negative TEs at specific electrodes and masker conditions (e.g., ID1 in the BP configuration; ID9, ID10, and ID11 in the optimized configurations), and in some cases this directionality varied within the same subject depending on stimulation site or masker. Given the small magnitude of these negative values and the test-retest variability inherent to psychophysical threshold measurements – estimated at 3.25 CU and 3.04 dB in a comparable EAS masking paradigm (Krüger et al., 2017) – these individual negative TEs cannot be distinguished from a threshold that is simply not significantly different from zero, and we do not interpret them as evidence that simultaneous acoustic stimulation can genuinely lower the electric detection threshold. We note, however, that mechanisms by which a masker can produce enhancement rather than suppression of a probe response have been described in the literature: Cheatham and Dallos (1982) showed in the cochlear microphonic (CM) that interference tones can either suppress or enhance the probe response depending on their frequency relative to the probe/recording site, attributing this to constructive or destructive vectorial summation of spatially distributed generator outputs. While such a mechanism offers a plausible explanation for why negative TEs can in principle occur in acoustic-electric masking paradigms, our study was not designed to test for enhancement at the individual level, and we therefore refrain from drawing conclusions about its presence in our data beyond nothing it as a descriptive observation for future, appropriately powered investigation.

In the present study we hypothesized that electric stimulation may reach the apex and interact with low-frequency stimulation. It is also possible that electrical stimulation at the cochlear base may activate the auditory nerve within the internal auditory canal, a phenomenon known as ectopic stimulation (e.g., Kalkman et al., 2016). This could also interact with low-frequency acoustic stimulation and thereby contribute to TE. 3D finite element method models coupled to physiological models of the auditory nerve to electric and acoustic stimulation (Kipping et al., 2024; Zhang et al., 2025) can be useful to understand the underlying mechanisms explaining acoustic masking of electric stimuli at the base or the RW.

### 4.5. Correlation between hearing loss and threshold elevation of acoustic maskers on electrical stimulation

The results indicate that TE during electrical stimulation delivered with the most apical electrode is partly related to residual hearing. Significant correlations were observed between TE and audiometric thresholds across frequencies from 125 Hz to 2000 Hz when TE was evaluated in CU. These findings indicate that better residual hearing is associated with stronger acoustic masking of electric stimulation delivered to the most apical electrode. In contrast, no correlation was observed when TE was expressed as %DR.

A similar pattern was observed for masking caused by a higher-frequency acoustic masker (e.g., 750 Hz) during apical electrical stimulation. Although the correlation did not reach statistical significance, the same tendency relating audiometric thresholds and TE was present only in CU but not in %DR. The lack of correlation when expressing TE in %DR may be explained by the large individual variability of DR across subjects. For the low-frequency masker (250 Hz), no correlation was found between TE and audiometric thresholds, either in CU or in %DR.

The results are consistent with findings by Imsiecke et al. (2019), who reported that subjects without measurable acoustic masking typically exhibit poorer PTA than those in whom masking is present. As in the present study, this relationship was most evident for apical stimulation sites, where EAS interactions are expected to be strongest.

In contrast, no significant correlation between residual hearing and TE was observed for stimulation delivered at basal electrodes or near the RW, irrespective of whether TE was expressed in CU or %DR. This indicates that, although acoustic masking can occur when electric stimulation is delivered at these stimulation sites, its magnitude is not systematically related to audiometric thresholds. Despite the limited consistency of correlations across maskers, assessing residual hearing may still offer diagnostic value, as masking effects could serve as an indicator of preserved cochlear function in individual cases. Analysis of current-flow based on lumped-parameter model did not reveal any significant relationship between amount of estimated current reaching the apex with TE. Within the scope of this study, current flow estimates therefore do not provide additional power for explaining inter-individual differences in TEs.

### 4.6. Clinical application

In clinical practice, low-frequency hearing is typically assessed using objective measures such as OAEs, ABRs, and ASSRs. While these techniques are well established for detecting hearing thresholds in infants and difficult-to-test populations, they show reduced reliability in the low-frequency range. OAEs are highly susceptible to noise and middle-ear effects, and low-frequency ABRs and ASSRs often require long averaging times and still suffer from poor signal-to-noise ratios. As a result, accurate diagnosis of low-frequency residual hearing in newborns remains challenging.

The present results suggest that acoustic masking of electrically evoked percepts could potentially serve as an alternative approach to assess low-frequency auditory function. Masking-based thresholds may offer a way to evaluate the functional status of low-frequency auditory nerve fibers, provided that electrical stimulation can selectively activate the relevant regions of the cochlea.

However, several limitations must be addressed before such an approach could be clinically implemented. First, electrical stimulation delivered through electrodes close to the RW may elicit SEs or non-auditory sensations, which reduce possibility of evoking sound sensations and consequently measuring acoustic masking. Second, reliable masking-based assessment requires that significant amount of electric stimulation effectively activates auditory pathways. In the present study we used CI electrodes that in principle are in close proximity to the RW but without control on their orientation and possible contact with cochlear structures. The use of transtympanic electrodes that can be placed in the promontory or close to the RW may be more effective for that purpose. Third, this study did not include formal cognitive testing of participants. Potential participants with documented cognitive impairments in their patient records were not invited to participate; however, no systematic cognitive screening was performed as part of the study protocol itself. Given the wide age range of the cohort (24–83 years, mean 59.2 years), undetected differences in cognitive function among included participants could have contributed to the considerable inter-subject variability observed in masking thresholds, as psychophysical detection and masking tasks generally place demands on attention, working memory, and response consistency. Future studies should therefore consider incorporating standardized cognitive screening to account for this potential confounding factor, particularly when extending this paradigm to pediatric or elderly populations.

Future work will therefore need to optimize electrode placement, stimulation parameters, and SE mitigation strategies to determine whether acoustic masking of electric stimulation delivered to close proximity of the RW can become a viable complementary tool for low-frequency hearing assessment in clinical populations such as newborns.

## 5. Conclusion

This study investigated psychoacoustic electric-acoustic masking when electrical stimulation was delivered through electrodes near the RW, analyzing the effects of different pulse rates, phase widths, and stimulation types. To evaluate the effectiveness of RW stimulation, trans-impedance measurements (TIMs) were used to analyze current flow, a loudness-scaling procedure was conducted to assess potential side effects (SEs), and acoustic masking was measured and compared for electrical stimulation delivered at the apex, the base, and near the RW.

Based on the recorded TIMs, a current-flow model was adapted to estimate current entering and leaving the cochlea. The model indicated that, during stimulation in the basal region, a larger proportion of current exits the cochlea, which can contribute to the occurrence of SEs. In contrast, apical intra-cochlear stimulation resulted in a current pattern that inherently directs more current toward apical, low-frequency regions. These findings suggest that stimulation delivered near the RW or at the base of the cochlea can potentially activate low-frequency nerve fibers more strongly, which may place them into a refractory state and thereby causing increase detection thresholds in the presence of an acoustic masker.

The results further confirmed that electrical stimulation through electrodes near the RW produces current flow into the cochlea and evokes sound sensations that increase in loudness with increasing current, despite the occurrence of SEs in some cases. The estimated electrical DR was greater for stimulation at the AP electrode compared to the BA electrode, with a significantly larger DR observed for BP pulses when using clinical pulse-rate and phase-duration settings.

Moreover, low-frequency acoustic stimulation produced significant masking of electrical stimulation delivered to the AP electrode in the BP configuration, whereas masking was less pronounced for stimulation delivered at the BA or RW electrodes. The results also revealed pronounced inter-individual variability in masking, indicating that higher-frequency acoustic stimulation can produce stronger masking effects when sufficient residual hearing is present.

Longer phase durations showed a tendency toward increased masking in optimized stimulation configurations at the individual level; however, these effects did not reach statistical significance at group level. A correlation analysis between residual hearing and TE caused by acoustic masking during electrical stimulation demonstrated a significant relationship when electric stimulation was delivered to the AP electrode, where masking effects were strongest. This correlation was most pronounced for higher-frequency or broader frequency maskers.

Overall, these results may be clinically relevant for assessing low-frequency hearing by means of acoustic masking of electric signals delivered to electrodes placed close to the RW, suggesting a potential application as a novel diagnostic approach.

## Data Availability Statement

The data that support the findings of this study are available upon reasonable request from the authors.

## Ethical Statement

This retrospective study analyzed previously published data. The original studies were carried out in accordance with the declaration of Helsinki principles and approved by the ethics committee of the Hannover Medical School (Hanover, Germany). The participants provided their written informed consent to participate in these studies.

## Acknowledgements

The authors would like to thank all study participants for their time and commitment. Funding for this research was provided by the European Research Council (ERC) under the European Union’s Horizon-ERC programme (Grant agreement No. 101044753) and by the Agencia Estatal de Investigación under the ATRAE project (Grant No. ATR2023-145064).

## Declaration of Generative AI and AI-assisted technologies in the writing process

During the preparation of this work, the author(s) used ChatGPT (OpenAI) and Claude (Anthropic) to support language contextualization, grammar, and spelling, and in part to assist with structuring the outline of the manuscript.

## Appendix 1

**Figure A 1.**
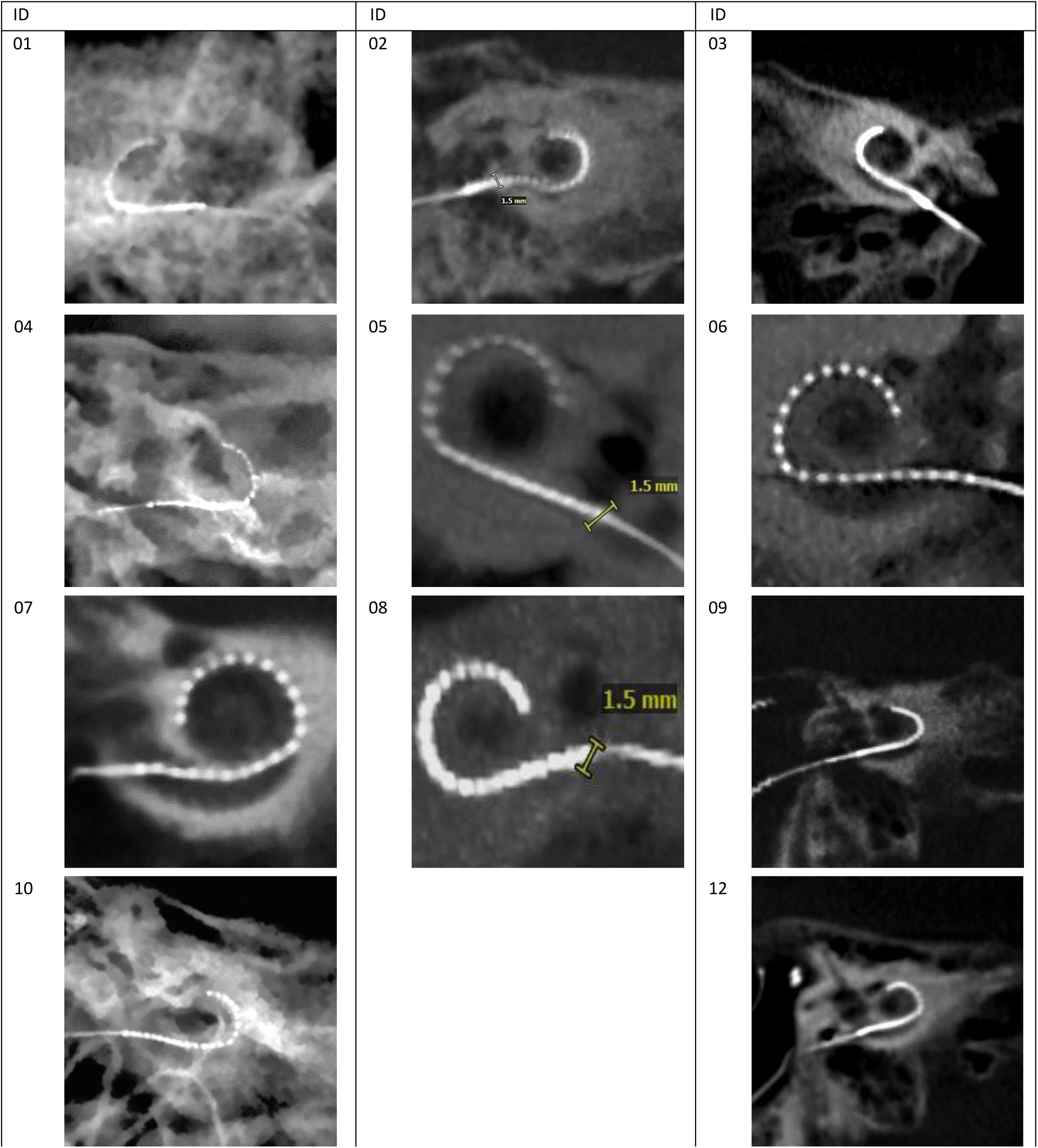
Illustration of cone beam computed tomography (CBCT) images for all subjects who participated, with the exception of Subject 11, whose CBCT is instead shown exemplarily in Figure 2 (main manuscript). For all subjects, the round window (RW) niche cross-section was determined from the images, together with the known inter-electrode distances of the implanted array (Hybrid-L, Slim Straight, or Slim 20; Cochlear Ltd.), following the protocol of Thormählen et al. (2024). Subjects 01 and 04 were imaged using X-ray rather than CBCT; for Subject 10, a processed X-ray image is shown in place of CBCT due to insufficient CBCT image quality. For these three subjects (01, 04, 10), RW electrode position was estimated based on the known electrode geometry rather than determined directly from the images, owing to the lower resolution of X-ray compared to CBCT. Image sections for Subjects 01, 09, 10, and 12 were additionally reprocessed to improve visibility of the round window and surrounding cochlear structures. In all subjects, an electrode contact was located at or near the RW niche; in Subject 07, the most basal electrode was inserted slightly deeper than in the other subjects, though still close to the RW niche. The subject ID is provided for identification purposes. The type of implanted electrode array can be found in Table 1.

